# Defective transcription of AAGAG satellite DNA causes sex-ratio meiotic drive in *Drosophila*

**DOI:** 10.1101/2025.01.10.632443

**Authors:** Tomohiro Kumon, Mami Nakamizo-Dojo, Amelie A Raz, Romain Lannes, Jaclyn M Fingerhut, Yukiko M Yamashita

## Abstract

Male germ cells have complex transcriptomes, with a large fraction of the genome being transcribed. This includes protein-coding genes (often not translated), non-coding DNA, and repetitive DNA, such as transposons and satellite DNA, which are normally silenced as heterochromatin. The significance of such widespread transcription remains unknown. Here, we show that a heterochromatin protein, HP2, is required for the transcription of AAGAG satellite DNA in *Drosophila* spermatocytes. HP2 depletion leads to abnormal retention of heterochromatin histone marks (H3K9me3) and spermatid death during sperm DNA packaging, leading to a model that transcription of AAGAG satellite DNA facilitates the remodeling of its heterochromatic nature in preparation for sperm DNA packaging. Strikingly, the severity of the spermatid death correlates with the amount of AAGAG satellite DNA carried by the spermatids, leading to preferential death of Y chromosome-containing spermatids over X-containing spermatids, and hence sex-ratio meiotic drive phenotype. We propose that widespread spermatocyte transcription may reflect the process of chromatin remodeling to allow sperm DNA packaging. We further propose that differential composition and amount of satellite DNA on chromosomes may underlie naturally occurring male meiotic drive.

## Introduction

In a broad range of organisms from *Drosophila melanogaster* to *Homo sapiens*, testis is known to have a high-complexity transcriptome, transcribing a large fraction of the genome. This includes many protein-coding genes (without necessarily being translated), non-coding DNAs, as well as repetitive sequences such as transposons and satellite DNA that are normally silenced as heterochromatin^1–8^. A few hypotheses have been proposed to explain such widespread transcription. For example, one model proposed that transcription-coupled DNA repair in male germ cells serves to lower mutation rates of protein-coding genes^9^(but see also ^10^), but it does not explain the significance of transcription of non-coding and repetitive sequences. Others proposed that non-coding transcripts function as lncRNA of yet unknown function during spermatogenesis, but many of them showed no phenotype upon deletion^2^. Yet another model proposed that transcription is a by-product of generally open chromatin in testis ^5^. Accordingly, there is no consensus model that explains widespread transcription of the testis to date.

Here using *Drosophila* spermatogenesis as a model (Fig. 1a), we investigate the role of satellite DNA transcription. Satellite DNA is non-coding tandem repetitive DNA, and constitutes approximately 30% of the *Drosophila* genome (Fig. 1b)^11–13^. We show that satellite DNA is highly transcribed in spermatocytes along with other transcripts, contributing to widespread transcription. We found that a heterochromatin binding protein, HP2 ^14–17^, which preferentially binds to AAGAG satellite DNA, is required for its transcription in spermatocytes. HP2 depletion led to subfertility due to defective sperm DNA packaging, associated with aberrant retention of heterochromatin histone mark H3K9me3 and defective incorporation of protamine, the sperm-specific DNA packaging protein^18, 19^. Strikingly, we found that defects in sperm DNA packaging upon HP2 depletion correlate with the abundance of AAGAG satellite DNA contained in the haploid spermatid: spermatids with the Y chromosome, which contains much more AAGAG satellite DNA than the X chromosome, were particularly sensitive to HP2 depletion, leading to skewed sex ratio in progeny. Moreover, spermatids containing a chromosome II variant with a minimal amount of AAGAG satellite DNA were more resistant to HP2 depletion, leading to a partial rescue of the HP2 depletion phenotype. Taken together, we propose that HP2-mediated AAGAG satellite DNA transcription facilitates the opening of AAGAG satellite DNA heterochromatin in preparation for sperm DNA packaging. We speculate that, in more general, widespread transcription broadly observed in male germ cells may reflect the process of chromatin remodeling during sperm DNA packaging. We further propose that the distinct composition of satellite DNA may have been exploited by meiotic drivers, leading to an evolutionary arms race between meiotic drivers and satellite DNA.

**Fig. 1:**
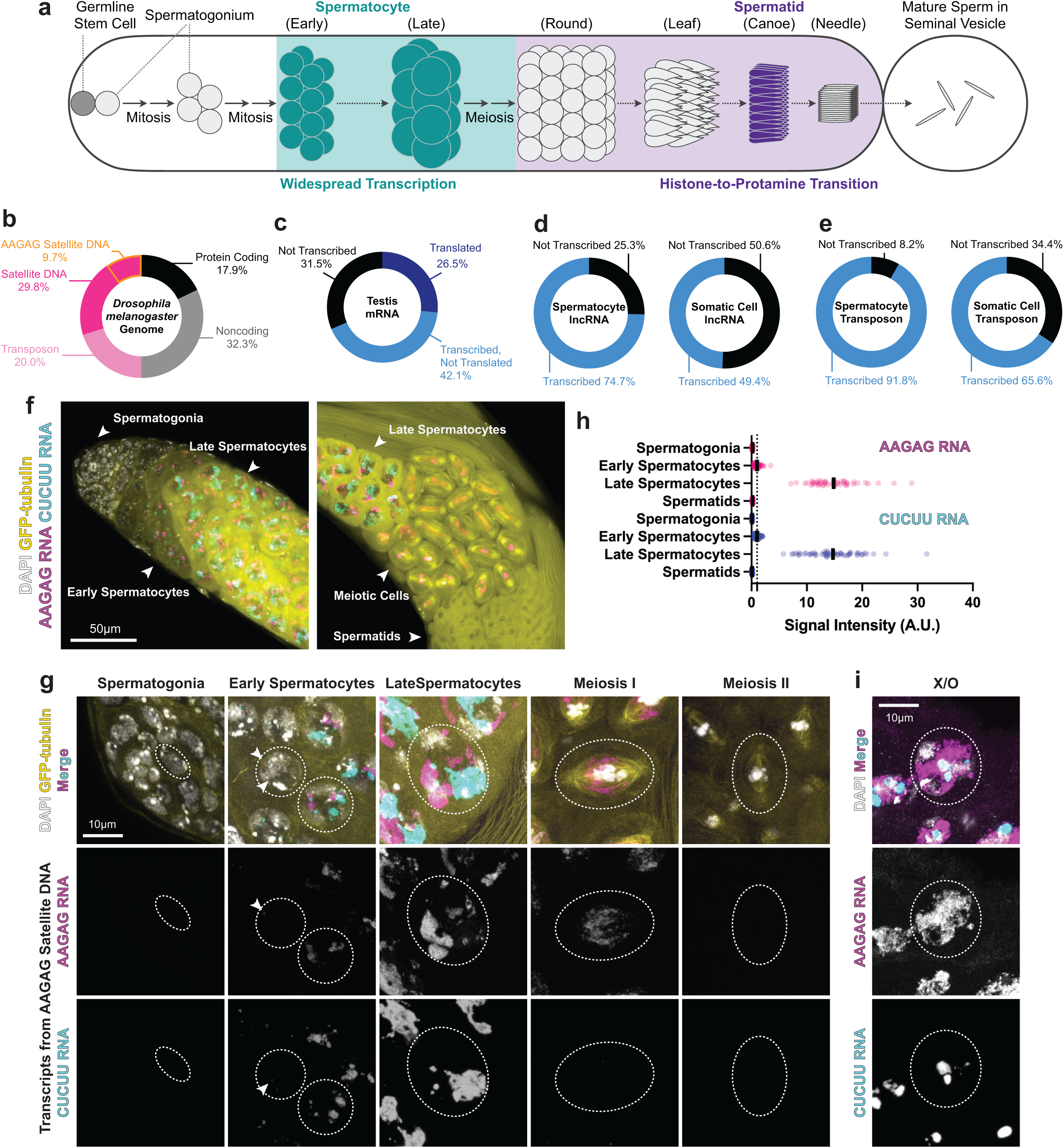
*Drosophila* spermatocytes exhibit widespread transcription. (a) Schematics of *Drosophila* spermatogenesis. Germline stem cells at the apical tip of the testis (left in the cartoon) produce differentiating spermatogonia, which divide four times before becoming spermatocytes. While spermatocytes grow in size, they exhibit widespread transcription, with many protein-coding and non-coding transcripts. After meiosis, spermatid nuclei undergo morphological changes through round, leaf, canoe, and needle stages, during which histone is replaced by protamine. Mature sperm are stored in seminal vesicles and prepared for ejaculation. (b) Composition of *Drosophila melanogaster* genome^13, 24, 52, 53^. AAGAG satellite DNA occupies approximately 10% of the genome. (c) Transcription of protein-coding genes (12,760 genes) in the *Drosophila* testis. Approximately 70% of total protein-coding genes are transcribed (30% translated (dark blue), and 40% not translated (blue)), while 30% are not transcribed (black). (d-e) Transcription of lncRNAs (1764 genes) and transposons (122 transposons) in spermatocytes vs. somatic cells (somatic epithelia surrounding each cyst). Note that the RNA sequencing data ^1^ were obtained by polyA selection, thus do not capture transcripts that are not polyadenylated such as satellite RNA. (f, g) RNA FISH of AAGAG/CUCUU satellite transcripts in male germ cells in various stages of differentiation (germline stem cells/spermatogonia, spermatocytes, meiotic cells). f) low magnification images capturing the course of germ cell differentiation. g) individual cells at various stages (signal intensities are not modified relative to each other). (h) Quantification of AAGAG/CUCUU satellite RNA at various stages of male germ cell differentiation based on FISH signal (see Methods). n = 50 cells were counted for each cell type. The black bar in each row represents the mean value. The dotted black line at 1 indicates the reference level. Signal intensities are shown as relative values normalized to the control (early spermatocytes), whose average value was set to 1 (as described in the Methods). Two biological replicates were performed. (i) RNA FISH of AAGAG/CUCUU satellite transcripts in the spermatocyte of X/O males lacking the Y chromosome, demonstrating these transcripts are not solely from the satellite DNA-rich Y chromosome.

## Results

### Satellite DNA is transcribed in *Drosophila* spermatocytes

We comprehensively analyzed the transcription in the *Drosophila* testis, utilizing multiple available data sets, and established the nature of widespread transcription. *Drosophila* genome is composed of ∼20% of protein-coding genes, ∼30% of satellite DNA, ∼20% of transposons, and ∼30% of other non-coding/non-repetitive DNA (Fig. 1b). Previous single-nucleus and single-cell RNA-seq data from *Drosophila* testis showed the widespread transcription of polyadenylated transcripts in the spermatocytes^1^. Reanalysis of these data^1^ further revealed that 69% of protein-coding genes are expressed in the testis (Fig. 1c), which is much higher than in somatic tissues^20^. Comparison of transcriptome^1^ and proteome^21^ further revealed that more than half of transcribed protein-coding genes (i.e., 42.1% of total protein-coding genes) are not translated to proteins (Fig. 1c). lncRNAs and transposons were more abundantly transcribed in spermatocytes than in the somatic epithelia of the testis (Fig. 1d, e).

Moreover, RNA fluorescence *in situ* hybridization (FISH) revealed marked upregulation of various satellite DNA transcription in spermatocytes, which cannot be captured by RNA sequencing that only identified polyadenylated RNAs^1^ (Fig. 1f, Supplementary Fig. 1). Most of satellite DNA found in *Drosophila melanogaster* genome^13, 22–24^ were found to be transcribed in spermatocytes, with many of them exhibiting the signal from both strands (Fig. 1f, Supplementary Fig. 1). Although a known source of satellite transcripts in the spermatocytes is satellite DNA embedded in the gigantic introns of several Y-linked genes ^24–27^, the large amount of satellite transcripts remained in the spermatocytes from X/O males that lack the Y chromosome (Fig. 1g, Supplementary Fig. 1), suggesting that satellite DNA from the X chromosome and autosomes are also transcribed. Taken together, these results establish the widespread nature of spermatocyte transcription in *Drosophila* spermatocytes, including protein-coding genes, non-coding RNA, and satellite DNA.

### HP2 is required for the transcription of AAGAG satellite DNA in spermatocytes

We found that heterochromatin protein 2 (*Su(var)2-HP2*, hereafter *HP2*), an HP1-interacting protein^14, 17^, is specifically required for the transcription of AAGAG satellite DNA in the spermatocytes. Depletion of HP2 from late spermatogonia/ spermatocytes using *bam-GAL4* driver (*bam-GAL4>UAS-HP2^HMS01699^*, hereafter *bam>HP2^RNAi^)* led to marked reduction of transcripts from AAGAG satellite DNA in spermatocytes: RNAs from both strands of AAGAG satellite DNA (i.e., AAGAG RNA and CUCUU RNA) were reduced in *bam>HP2^RNAi^*, with a more severe reduction of AAGAG RNA than CUCUU RNA (Fig. 2a, b). Expression of RNAi-resistant HP2 construct rescued transcription of AAGAG satellite DNA (both strands) in *bam>HP2^RNAi^* (Supplementary Fig. S2), demonstrating that reduction in AAGAG/CUCUU RNA is indeed caused by HP2 depletion. *bam>HP2^RNAi^* had little effect on the transcription of other satellite DNA (Supplementary Fig. S3), suggesting that HP2 regulates satellite DNA transcription in a sequence-specific manner. These results demonstrate that HP2 is required for AAGAG/CUCUU transcription in spermatocytes.

**Fig. 2:**
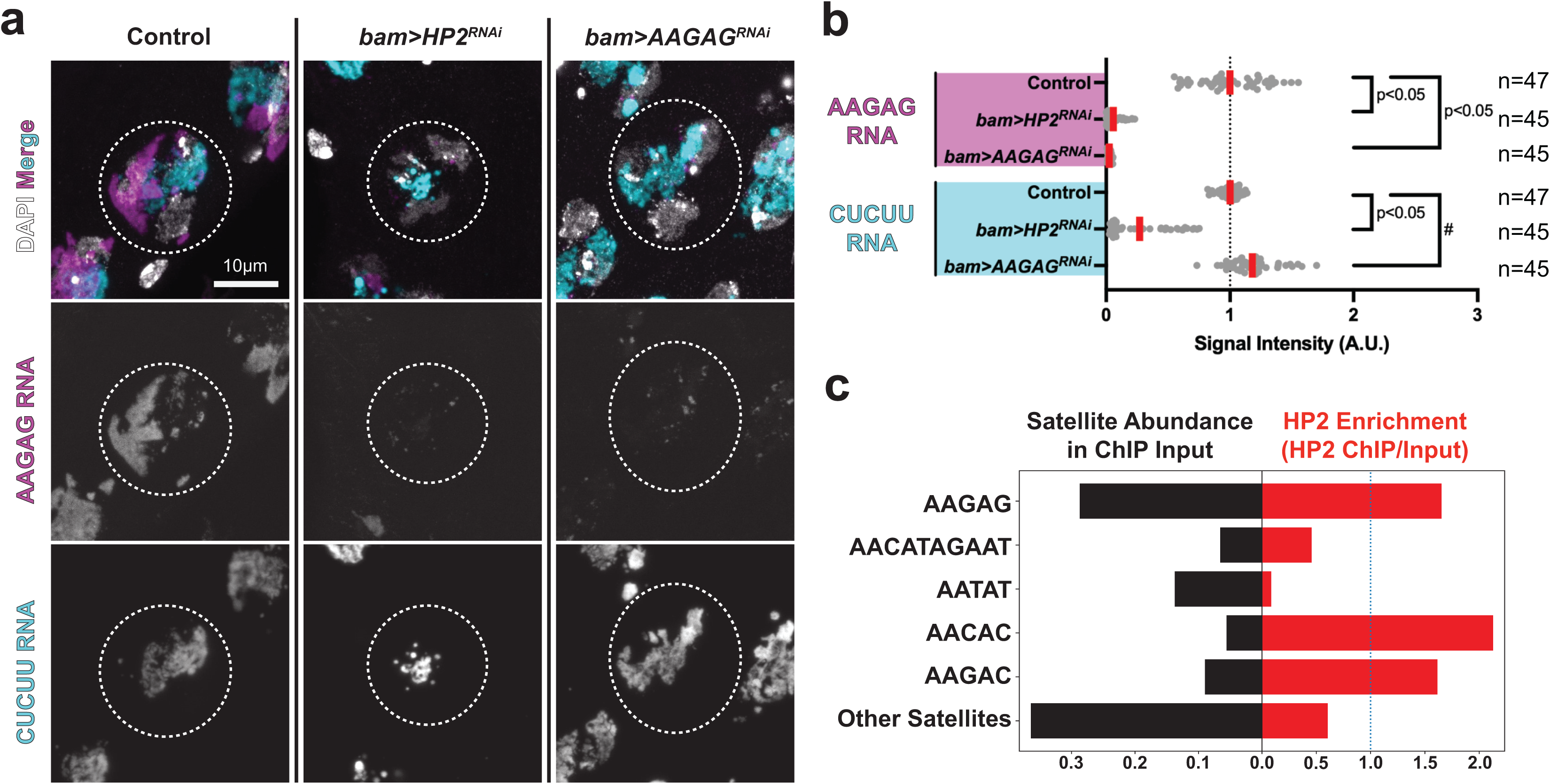
HP2 is required for the transcription of AAGAG satellite DNA in spermatocytes. (a) RNA FISH of AAGAG/CUCUU satellite transcripts in control, *bam>HP2^RNAi^* and *bam>AAGAG^RNAi^* spermatocytes. Spermatocyte nuclei are indicated by dotted circle. (b) Relative signal intensities of AAGAG/CUCUU transcripts in control, *bam>HP2^RNAi^* and *bam>AAGAG^RNAi^* spermatocytes (n = number of spermatocytes examined). Red line: mean. The *p*-value is calculated by an unpaired t-test (two-tailed). #: CUCUU RNA was slightly but significantly (p<0.05) upregulated in *bam>AAGAG^RNAi^*. Each dot represents the RNA FISH intensity in a single spermatocyte (Method). Exacat *p*-values are provided in the Source Data file. (c) HP2 enrichment on satellite DNA (right) and the abundance of satellite DNA in the *Drosophila melanogaster* genome calculated from the ChIP input (left). HP2 ChIP-seq data (modENCODE ID: 5593)^28^ were analyzed using the k-Seek method ^36^ to quantify simple satellite repeat abundance from short-read sequencing (Method).

Analysis of HP2 ChIP-seq data on the modENCODE database^28^ showed that HP2 binds preferentially to AAGAG satellite DNA (Fig. 2c), implying that HP2 may regulate the transcription of AAGAG satellite DNA via direct binding. Moreover, we found that HP2’s localization to chromatin is dependent on AAGAG satellite DNA (Supplementary Fig. S4). However, given that HP2 expression decreased at the time of strongest AAGAG/CUCUU transcription in spermatocytes (Supplementary Fig. S4a), it is possible that HP2’s role in AAGAG/CUCUU transcription can be indirect. If it is direct, HP2’s requirement in AAGAG/CUCUU transcription may represent an example of heterochromatin-dependent transcription^29, 30^.

It was previously shown that AAGAG RNA can be depleted by RNAi (*bam-GAL4>UAS-AAGAG^shRNA^*, hereafter *bam>AAGAG^RNAi^*)^3, 31^. Although the mechanism of nuclear RNA degradation by RNAi is not fully established, *bam>AAGAG^RNAi^* was previously used to eliminate AAGAG RNAi from the spermatocytes^3^. We confirmed that *bam>AAGAG^RNAi^* specifically depleted AAGAG RNA without affecting the opposite strand, CUCUU RNA (Fig. 2a, b) as previously shown^3^. These results suggest that *bam>HP2^RNAi^* interferes with the transcription of AAGAG satellite DNA, whereas *bam>AAGAG^RNAi^* cleaves AAGAG RNA after it is transcribed. Hereafter, we analyzed both *bam>HP2^RNAi^* and *bam>AAGAG^RNAi^* because their comparison provided insights into the function of AAGAG transcription vs. AAGAG transcript. In conclusion, HP2 is required for the transcription of AAGAG satellite DNA.

### HP2 depletion leads to subfertility due to sperm DNA packaging defects

We found that *bam>HP2^RNAi^* led to a reduction in fertility, whereas *bam>AAGAG^RNAi^* led to complete sterility, as shown previously^3^ (Fig. 3a). The majority of *bam>HP2^RNAi^* animals still had mature sperm in their seminal vesicles, corresponding to their residual fertility, whereas *bam>AAGAG^RNAi^* animals had no sperm in their seminal vesicles (Fig. 3b). Based on the sterility caused by *bam>AAGAG^RNAi^*, a previous study proposed that AAGAG RNA plays an important role in the process of spermatogenesis. They reported that *bam>AAGAG^RNAi^* led to late spermatogenesis phenotypes, such as nuclear morphology defects and sperm bundling defects^3^. They further observed that *bam>AAGAG^RNAi^* led to defective protamine incorporation into sperm nuclei, a critical process for sperm DNA packaging. However, we found that protamine was mostly normally incorporated in *bam>AAGAG^RNAi^* testis (Supplementary Fig. S5). We noted that the disruption of sperm nuclei bundling made it challenging to correctly identify the differentiation stages of sperm nuclei, potentially leading to the interpretation of defective protamine incorporation in the previous study^3^. RT-qPCR showed that *bam>AAGAG^RNAi^* led to marked downregulation of a subset of Y-linked fertility genes (e.g., *ORY* and *kl-2*) (Fig. 3c). *kl-2* encodes a subunit of axonemal dynein, whose depletion is known to lead to the disruption of sperm nuclei bundling^27^, similar to the phenotype of *bam>AAGAG^RNAi^*. *ORY* and *kl-2* genes are known to contain large stretches of AAGAG satellite DNA in their gigantic introns (Supplementary Fig. S6)^24, 26^. RT-qPCR further revealed a striking reduction of *ORY* transcripts across AAGAG-containing gigantic introns in *bam>AAGAG^RNAi^* animals (Fig. 3d). RNA FISH further confirmed the reduction of *ORY* mRNA in spermatocytes in *bam>AAGAG^RNAi^* animals (Figure 3e). Moreover, *Pzl*, an autosomal gene that contains gigantic introns with assembly gaps^32^ that likely contain AAGAG satellite DNA (Supplementary Fig. S6), was also downregulated in *bam>AAGAG^RNAi^* (Fig. 3c). These results suggest that *bam>AAGAG^RNAi^* leads to cleavage of AAGAG-containing RNA. Because *ORY* and *kl-2* are required for male fertility^33–35^, these results suggest that the sterility of *bam>AAGAG^RNAi^* is caused by RNAi-mediated knockdown of these genes that contain AAGAG in their introns.

**Fig. 3:**
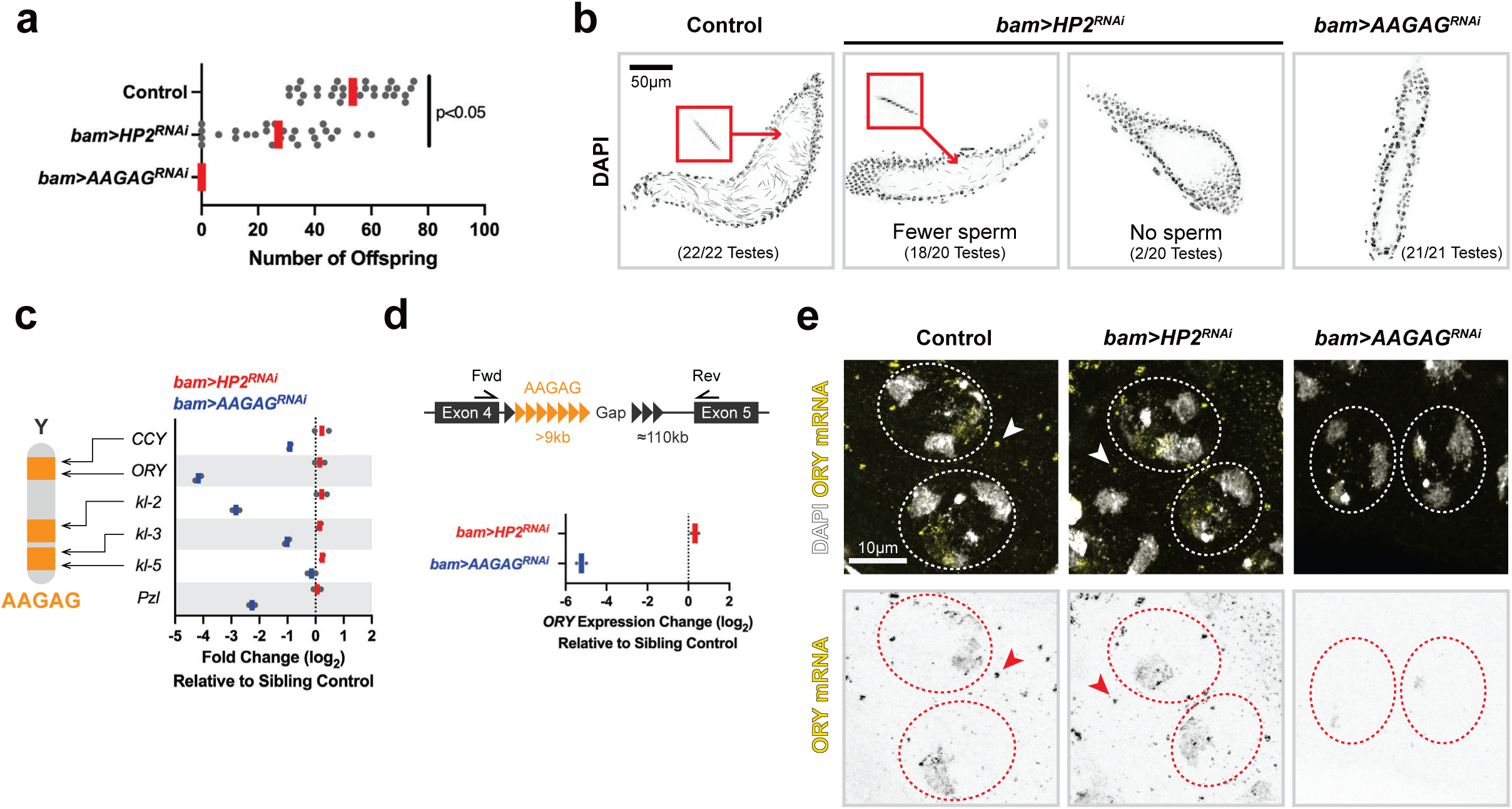
HP2 depletion leads to subfertility. (a) Fertility assay of wild type, *bam>HP2^RNAi^* and *bam>AAGAG^RNAi^* males. Each dot represents the number of offspring from a single mating pair (n = 29 mating pairs for control, n = 27 for *bam>HP2^RNAi^* and *bam>AAGAG^RNAi^*); red line, mean. The exact *p*-value is 0.000000071, which was calculated by an unpaired t-test (two-tailed). (b) Seminal vesicles of wild type, *bam>HP2^RNAi^* and *bam>AAGAG^RNAi^* males stained with DAPI. Only the seminal vesicle was cropped for clarity using Fiji/ImageJ. An example of sperm nucleus is marked with red. (c) Cytological map of AAGAG satellite DNA and fertility genes on the Y chromosome^13, 22, 24^. *Pzl* is an autosomal gene with large intronic AAGAG satellite DNA (Supplementary Figure S4). RT-qPCR using the primers to amplify the last exon of indicated genes in *bam>HP2^RNAi^* and *bam>AAGAG^RNAi^* compared to control with technical duplicates for each genotype. (d) RT-qPCR using the primers to amplify exon-exon junction across AAGAG satellite DNA-containing introns of *ORY* gene with technical duplicates for each genotype. (e) RNA FISH of *ORY* mRNA in spermatocytes in control, *bam>HP2^RNAi^* and *bam>AAGAG^RNAi^*. Cytoplasmic *ORY* mRNA granules (arrowheads) are formed in control and *bam>HP2^RNAi^*, but not in *bam>AAGAG^RNAi^*.

In contrast to *bam>AAGAG^RNAi^*, *bam>HP2^RNAi^* did not affect the expression of genes that contain gigantic introns with AAGAG satellite DNA (Fig. 3c, d, e), suggesting that *bam>HP2^RNAi^* impacts spermatogenesis through a mechanism distinct from *bam>AAGAG^RNAi^*. We found that *bam>HP2^RNAi^* led to sperm DNA packaging defects, explaining their subfertility. Although germ cell development appeared mostly normal, leading to normal meiosis (Supplementary Fig. S7), *bam>HP2^RNAi^* led to unique cytological phenotypes during spermatid differentiation stage. In wild type animals, post-meiotic germ cells undergo stereotypical morphological changes, leading to the formation of mature sperm: a cyst of 64 spermatids, resulting from four mitotic and two meiotic divisions, develop in synchrony to undergo sperm DNA packaging and axoneme elongation (Fig. 1a). Sperm DNA is packaged by protamine, sperm-specific DNA packaging proteins that replace histone-based chromatin, leading to highly compacted DNA (Fig. 4a)^18, 19^. We found that *bam>HP2^RNAi^* led to defective sperm DNA compaction during the last stages of sperm development (Fig. 4b): In some cases (Fig. 4b, ii), a subset of spermatid nuclei within a cyst lost their typical ‘needle’ morphology (Fig. 4b, i) and become completely rounded-up, while other nuclei exhibited normal needle morphology. In other cases, all spermatids within a cyst (Fig. 4b, iii) exhibited defective sperm nuclear morphology. Expression of RNAi-resistant HP2 rescued sperm DNA compaction defect in *bam>HP2^RNAi^* (Fig. 4b, iv), demonstrating that sperm DNA compaction defect is indeed due to HP2 depletion.

**Fig. 4:**
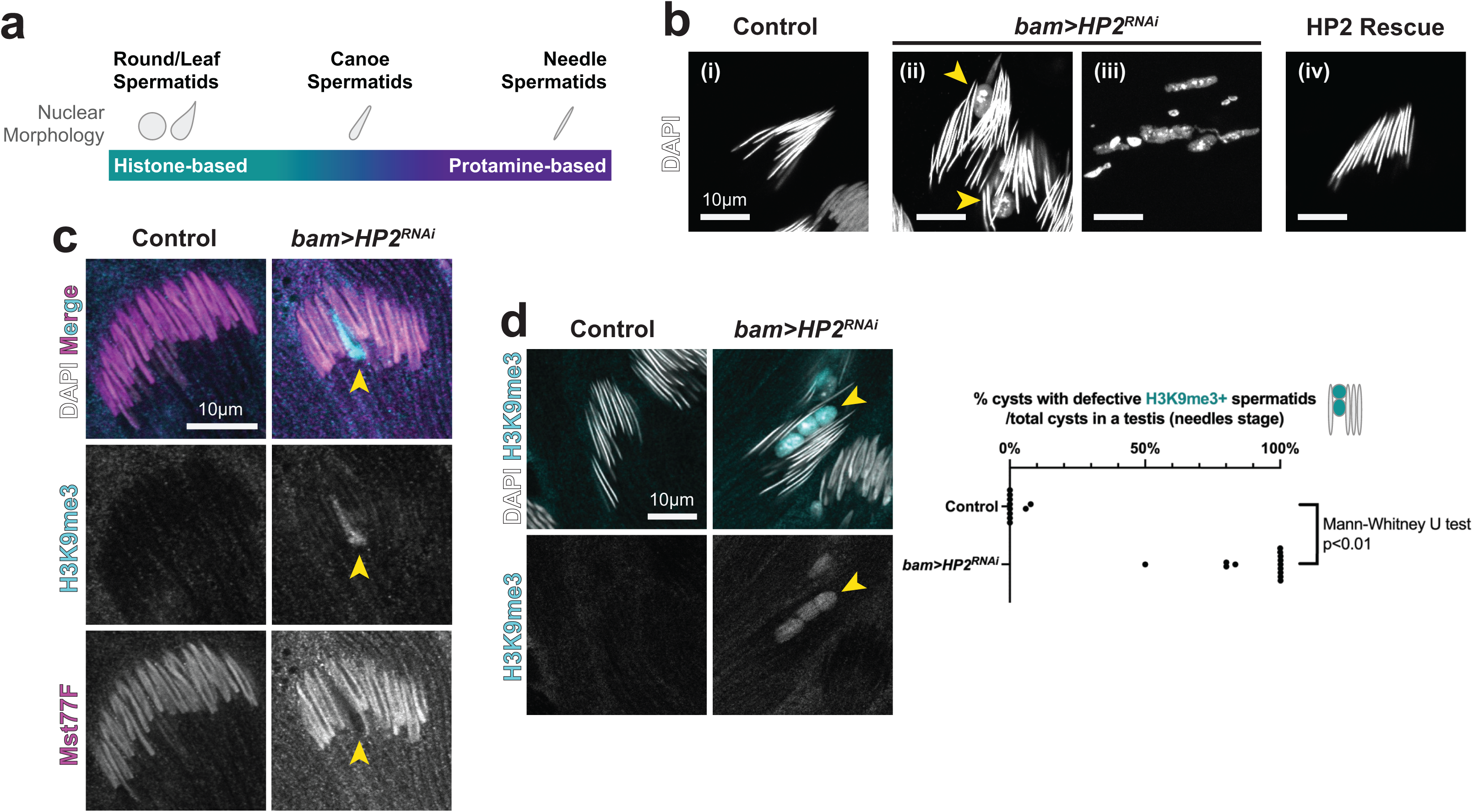
HP2 depletion leads to defective histone-to-protamine transition in spermatids. (a) Schematics of sperm development and histone-to-protamine transition. Histone-to-protamine transition occurs in the canoe stage spermatids. (b) Needle stage spermatids in control and *bam>HP2^RNAi^* animals, and *bam>HP2^RNAi^* animals with HP2 rescue construct (RNAi-resistant HP2-S transgene) stained with DAPI. Spermatid nuclei with defective sperm DNA compaction is indicated by arrowheads in ii. In some cysts (iii), all spermatids within a cyst failed to compact DNA. (c) Immunofluorescence staining of H3K9me3 and Mst77F in canoe spermatids from control and *bam>HP2^RNAi^* animals. Arrowheads indicate a spermatid nucleus that failed to undergo histone-to-protamine transition (n=15 testes for each genotype). (d) Immunofluorescence staining of H3K9me3 in needle spermatids from control and *bam>HP2^RNAi^* animals. Arrowhead indicates spermatid nuclei in *bam>HP2^RNAi^* animals that failed to undergo sperm DNA compaction and retained the H3K9me3 histone mark. Frequencies of cysts with H3K9me3 histone mark in needle stage spermatid in control and *bam>HP2^RNAi^* animals is shown. (n = 10 testes for control, n = 13 testes for *bam>HP2^RNAi^*). The exact *p*-value with the Mann-Whitney U test (two-tailed) is 0.00000175.

Although the morphology of earlier “canoe” stage spermatids appeared mostly normal in *bam>HP2^RNAi^* animals, we found that the defects began in these earlier stages. Canoe stage spermatid is characterized by the removal of histone and incorporation of protamine, such as Mst77F^13^. In control animals, sperm DNA compaction is accompanied by sequential events of removing heterochromatin mark H3K9me3, removing histone H3.3 (Supplementary Fig. S8a), then incorporating protamine such as Mst77F (Fig. 4c, control). However, in *bam>HP2^RNAi^* animals, a subset of spermatids exhibited a failure in the removal of H3K9me3 and deposition of Mst77F, whereas other nuclei in the same cyst have removed H3K9me3 and deposited Mst77F (Fig. 4c, *bam>HP2^RNAi^*). In wild type, protamine incorporation occurs in synchrony within the cyst, and all nuclei in a cyst are either Mst77F-positive or -negative (Fig. 4c, control). In contrast, the spermatids cyst from *bam>HP2^RNAi^* animals contained a mixture of Mst77F-positive and - negative nuclei, demonstrating that a subset of nuclei fails to incorporate Mst77F in a timely manner. Moreover, when the cyst progressed to the needle spermatid stage, the defective nuclei in *bam>HP2^RNAi^* that failed to compact their DNA still retained H3K9me3 (Fig. 4d). Similar to the retention of H3K9me3, H3.3 histone was also retained in the defective spermatids in *bam>HP2^RNAi^*, whereas another histone mark, AcH4, did not exhibit detectable difference between control and *bam>HP2^RNAi^* (Supplementary Fig. S8b). Taken together, these results suggest that retention of H3K9me3 is associated with the failure in protamine incorporation, which in turn leads to ultimate failure in sperm DNA compaction, in *bam>HP2^RNAi^*.

### HP2 depletion preferentially harms Y-chromosome-containing spermatids, leading to sex ratio meiotic drive

Unexpectedly, we found that *bam>HP2^RNAi^* preferentially impacts sperm DNA compaction of Y-chromosome-bearing spermatids, leading to a sex ratio meiotic drive phenotype. While counting the number of offspring for the fertility assay (Fig. 3a), we noticed that *bam>HP2^RNAi^* males produce significantly more daughters than sons (Fig. 5a), suggesting that sperm DNA compaction defects may be biased toward Y chromosome-containing spermatids. Indeed, by DNA FISH using X- and Y-specific probes, we found that defective nuclei in *bam>HP2^RNAi^* predominantly contained the Y chromosome (Fig. 5b), demonstrating that the Y chromosome is more sensitive to HP2 depletion than the X chromosome. Seeking for the difference(s) between X and Y chromosomes that may explain Y-biased sensitivity to HP2 depletion, we realized that the Y chromosome contains much more AAGAG satellite DNA than the X chromosome^13, 22^ (Fig. 5c). Based on this correlation between the amount of AAGAG on the chromosome and the tendency to fail in sperm DNA compaction, we hypothesized that HP2-dependent transcription of AAGAG satellite DNA prepares for protamine incorporation by remodeling heterochromatin state of AAGAG satellite DNA (Fig. 5d). Accordingly, the dependence of a chromosome on HP2 to remodel their AAGAG satellite DNA heterochromatin correlates with the amount of AAGAG satellite DNA on it. This can explain why X chromosome-containing spermatids are less impacted by HP2 depletion, because the X chromosome has less AAGAG satellite DNA. Based on these results, we propose that satellite DNA transcription, instead of its RNA product, is important to prepare for histone-to-protamine transition.

**Fig. 5:**
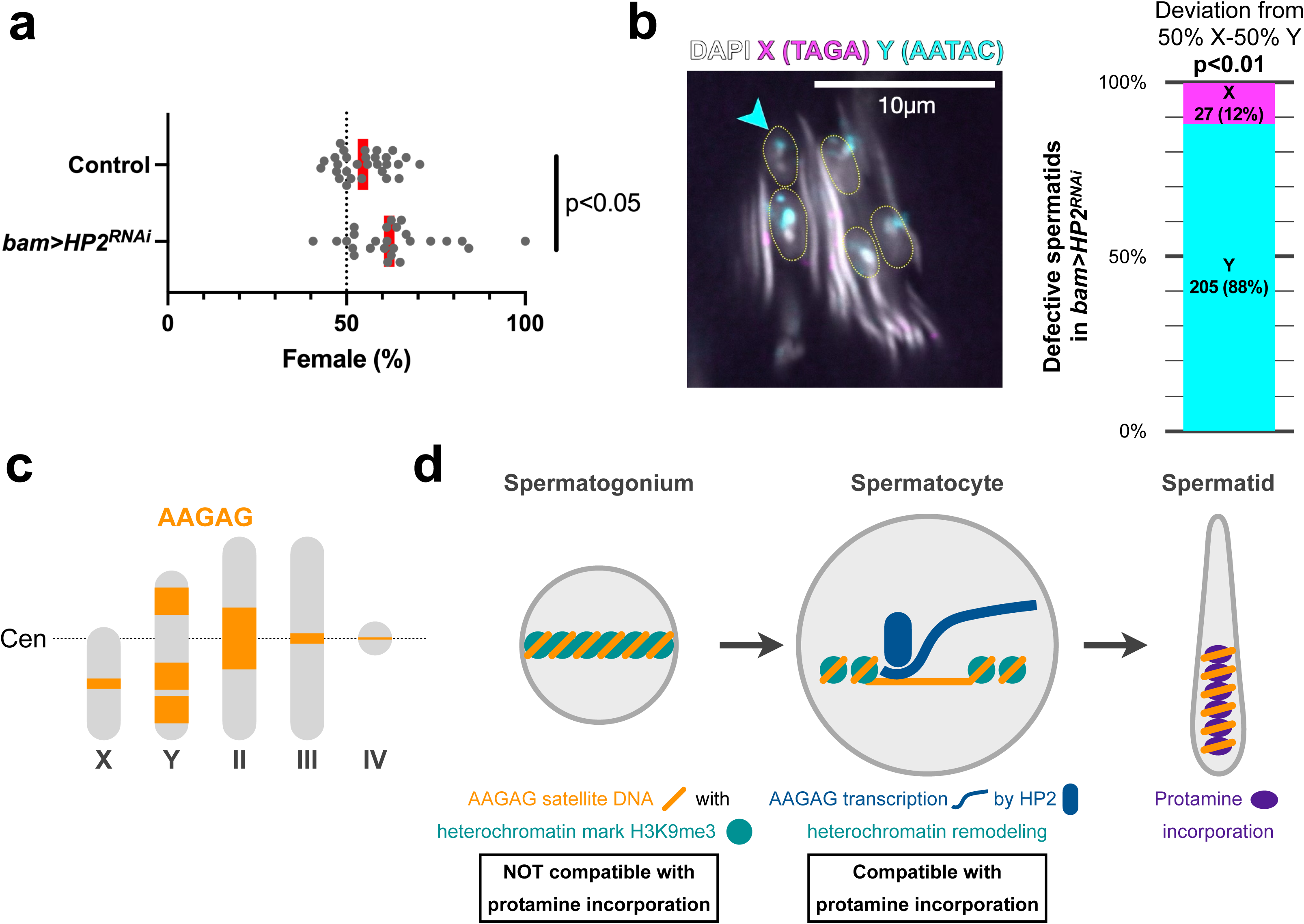
HP2 depletion preferentially harms Y-chromosome-containing spermatids, leading to sex ratio meiotic drive. (a) Sex ratio of the offspring of control vs. *bam>HP2^RNAi^* males. Each dot represents the sex ratio of offspring from a single mating pair (n = 29 for control, n = 26 for *bam>HP2^RNAi^*); red line, geometric mean. The exact *p*-value is 0.0052, which was calculated by an unpaired t-test (two-tailed). (b) DNA FISH on needle stage spermatids in *bam>HP2^RNAi^* spermatids using X- and Y-specific satellite DNA X: TAGA satellite DNA. Y: AATAC satellite DNA. The frequency of defective spermatids with X vs. Y chromosome is shown (n=232 spermatids from 14 testes). The exact *p*-value is 0.0001, which was calculated by the chi-square test for goodness of fit for deviations from 50% X and 50% Y. (c) Schematics of *Drosophila melanogaster* karyotype with AAGAG satellite DNA location (orange block, size is proportional to the estimate^13^). Large AAGAG satellite blocks are found in intergenic regions of the Y chromosome and pericentromeric region of the chromosome II. (d) Model for the functional significance of transcription.

### The amount of AAGAG satellite DNA determines sperm DNA compaction defects caused by HP2 depletion

We found that the reduction in AAGAG satellite DNA abundance can rescue sperm DNA compaction defect observed in *bam>HP2^RNAi^*, further supporting our hypothesis that the remodeling of satellite DNA via transcription is critical for sperm DNA packaging (Fig. 5d). In addition to the Y chromosome, chromosome II contains a considerable amount of AAGAG satellite DNA in its pericentromeric heterochromatin (Fig. 5c)^13, 22^, likely contributing to spermatid DNA compaction defects in *bam>HP2^RNAi^*. This may also explain why sometimes all spermatids within a cyst fail to undergo sperm DNA compaction defects (Fig. 4b, iii), because all spermatids contain a copy of chromosome II.

Consistent with this notion, we found that natural chromosome II variants with minimal AAGAG satellite DNA (hereafter II^ΔAAGAG^, ‘Ithaca 06’ and ‘Ithaca16’ strains)^36^ (Supplementary Fig. S9) can rescue sperm DNA compaction defects caused by the depletion of *HP2*. When this chromosome II^ΔAAGAG^ was placed in the background of *bam>HP2^RNAi^* (*y^+^w^+^/Y; UAS-HP2^HMS01699^ / ΔAAGAG; bam-GAL4/+,* hereafter X/Y; II^+^/II^ΔAAGAG^ *bam>HP2^RNAi^*), we observed a considerable reduction in the frequency of defective spermatids that failed sperm DNA compaction (Fig. 6a, b), suggesting that the abundance of AAGAG satellite DNA correlates with the severity of sperm DNA compaction defects due to HP2 depletion.

**Fig. 6:**
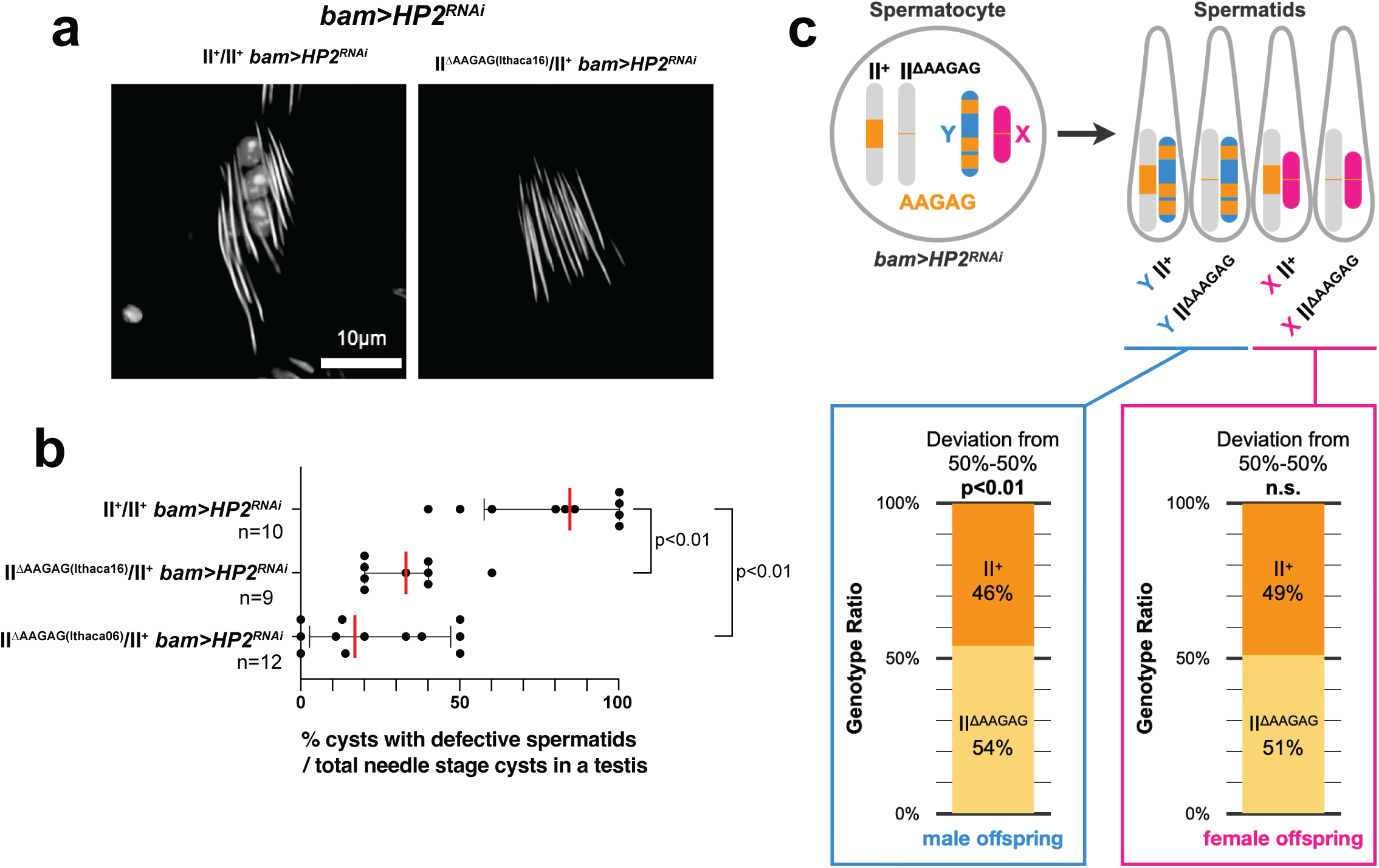
The amount of AAGAG satellite DNA determines sperm DNA compaction defects caused by HP2 depletion. (a) Needle stage spermatids stained with DAPI from *bam>HP2^RNAi^* animals with animals with indicated chromosome II genotypes (II^+^/II^+^ and II^ΔAAGAG^/II^+^). (b) The frequency of cysts with defective spermatids (at needle stage) was scored for the indicated genotypes. Each data point represents an individual testis. The red bar indicates the median, and the black bar indicates the interquartile range. The p-value was calculated by the Mann-Whitney U test (two-tailed). n = number of testes examined. The exact *p*-values were 0.0003 between control and ithaca16, 0.0001 between control and ithaca06. (c) Genotype of spermatids generated from X/Y; II+/II^ΔAAGAG^ *bam>HP2^RNAi^* spermatocytes, and their frequency represented in the offspring. n=1145 male offspring and n=296 female offspring were scored. The exact *p*-values were 0.00235 (in males) 0.727 (in females), calculated by the chi-square test for goodness of fit for deviations from 50% II^+^ and 50% II^ΔAAGAG^.

To further test our hypothesis that AAGAG satellite DNA amount determines the sensitivity of spermatids to HP2 depletion, we examined whether spermatids containing II^ΔAAGAG^ survived better than those with II^+^ in *bam>HP2^RNAi^*. Specifically, we analyzed the genotype of the offspring of X/Y; II^+^/II^ΔAAGAG^ *bam>HP2^RNAi^* males. From diploid X/Y; II^+^/II^ΔAAGAG^ spermatocytes, there are four possible genotypes of the haploid sperm: X II^+^, X II^ΔAAGAG^, Y II^+^, and Y II^ΔAAGAG^ (Fig. 6c). Based on the offspring genotype, we deduced the genotype of survived sperm. In male offspring, where II^+^ and II^ΔAAGAG^ can easily be scored based on the body color (Method), we observed a small but significant overrepresentation of II^ΔAAGAG^ over II^+^ (Fig. 6c), suggesting that spermatid containing Y II^ΔAAGAG^ indeed survived better than Y II^+^. This implies that AAGAG satellite DNA on chromosome II worsens sperm DNA compaction defects in Y-containing spermatids.

In female offspring, II^+^ and II^ΔAAGAG^ cannot be easily distinguished by visible phenotypic markers. Thus, a randomly-chosen subset of female offspring was subjected to a PCR-based assay to distinguish between II^+^ and II^ΔAAGAG^ (Method), which showed no significant difference in the survival of X II^+^ and X II^ΔAAGAG^ (Fig. 6c). This is not surprising considering that *bam>HP2^RNAi^* males with a regular amount of AAGAG (II^+^) sired female-biased offspring (Fig. 5a), suggesting that X-containing spermatids can tolerate the amount of AAGAG on the second chromosome. These results may reflect the dose-sensitive nature, where HP2-depleted cells can tolerate only a certain amount of AAGAG satellite DNA (see Discussion): X-containing spermatids may be able to tolerate AAGAG satellite DNA on the chromosome II, because they contain only a small amount of AAGAG satellite DNA on the X chromosome. In contrast, Y-containing spermatids, which carries a large amount of AAGAG satellite DNA, cannot tolerate AAGAG satellite DNA on the chromosome II under HP2 depletion. Accordingly, sperm DNA compaction defects of Y-containing spermatids in *bam>HP2^RNAi^* can be rescued by reducing the dose of AAGAG satellite DNA on the chromosome II.

These results suggest that the abundance of AAGAG satellite DNA is a determinant that renders spermatid sensitive to the loss of HP2. The fact that the reduction of AAGAG satellite DNA markedly rescues the phenotype of *bam>HP2^RNAi^* implies that AAGAG satellite DNA is the major target of HP2, although HP2 may also regulate other heterochromatic loci as well. Additionally, the rescue of the phenotype by reducing the amount of AAGAG satellite DNA implies that AAGAG RNA is not essential for spermatid development. Taken together, we propose that HP2 promotes transcription of AAGAG satellite DNA in spermatocytes, facilitating the remodeling of heterochromatin in preparation of germline DNA for histone-to-protamine transition.

## Discussion

Through the study of satellite DNA transcription in *Drosophila* spermatocytes, we propose that the widespread transcription in male germline reflects the process of chromatin remodeling in preparation for histone-to-protamine transition. According to this model, transcription, but not transcripts, is important for male germ cell development. Although it is challenging to distinguish between the function of transcription to remodel the chromatin and that to generate functional transcripts, several lines of evidence favor the model that transcription, but not resultant transcripts, is important. First, the expression of AAGAG did not rescue the *bam>HP2^RNAi^* phenotype (Supplementary Figure S10), suggesting that the lack of AAGAG transcript is not the cause of the *bam>HP2^RNAi^* phenotype. Furthermore, the rescue of the *bam>HP2^RNAi^* phenotype by the introduction of chromosome II with little AAGAG satellite DNA (II^ΔAAGAG^) (Fig. 6) provides critical support for this notion. If AAGAG RNA has an important function for spermatid development, it would be expected that deletion of AAGAG satellite DNA in II^ΔAAGAG^ worsens the phenotype of *bam>HP2^RNAi^* because both conditions reduce AAGAG RNA. Instead, II^ΔAAGAG^ rescued *bam>HP2^RNAi^* phenotype, implying that AAGAG RNA is dispensable for spermatid development. The role of AAGAG RNA indicated by the previous study by the use of *AAGAG^RNAi^* ^3^ likely reflects the function of fertility genes that contain AAGAG satellite DNA in their introns. Instead of AAGAG RNA serving a function, we propose that AAGAG RNA is produced as a consequence of transcription-dependent chromatin remodeling of heterochromatin, which is required for subsequent sperm DNA packaging by protamine. In this model, satellite DNA’s heterochromatic nature is a “burden” for spermatid development, impeding protamine-mediated sperm DNA compaction process if left unopened.

It is important to note that defective spermatid development in *HP2^RNAi^* is sensitive to the dose of total AAGAG carried in the haploid spermatids. We speculate that each spermatid may have a threshold of the unremodeled chromatin it can tolerate (e.g. the amount of unremoved histones, and/or the amount of the chromatin that is not packaged by protamine), and the spermatids that have an above-threshold unremodeled chromatin may be fated for death. Indeed, the presence of a checkpoint mechanism that monitors the quality of sperm DNA has been proposed to explain the cellular phenotype of sperm-killing meiotic drivers, such as *SD* ^37,38^. Interestingly, the cytological phenotype of *HP2^RNAi^* is reminiscent of the *SD* phenotype. We speculate that the AAGAG dose-sensitive nature of the *HP2^RNAi^* phenotype may be due to such a checkpoint mechanism.

HP2-mediated transcription of AAGAG satellite DNA likely represents just one example of many transcription regulators in the testis that reorganize chromatin for protamine incorporation. Among many heterochromatic satellite DNA blocks in *Drosophila* genome, HP2 appears to regulate mainly the transcription of AAGAG, whereas other satellite DNA such as AACATAGAAT, AATAT, AACAC, and AAGAC were not detectably impacted. Therefore, we speculate that there are other factors that serve the same function as HP2 but for different satellite DNA. By extending this model, we further propose that the widespread transcription of the testis, encompassing various genomic elements such as protein-coding (but not translated) genes and non-coding sequences, is the result of the process that remodels germ cell chromatin to allow histone-to-protamine transition. We predict that a panel of transcription factors together contribute to transcription of a large fraction of the genome in male germline. Discovery of such factors awaits future investigations.

Importantly, our study unexpectedly revealed a condition that caused sex-ratio meiotic drive. Because the Y chromosome contains more AAGAG satellite DNA, it was more sensitive to HP2 depletion than the X chromosome, causing HP2-depleted males to sire female-biased progeny. Our results may provide insights into the molecular mechanisms of evolution of satellite DNA and their binding proteins, eventually contributing to incompatibility between species. For example, if a chromosome that happens to have fewer copies of certain satellite DNA compared to other alleles in the population, such a chromosome can become a selfish meiotic driver by acquiring a mutation (e.g., dominant negative, or loss of function if haplo-insufficient) in the gene(s) that mediate transcription/remodeling of the target satellite DNA (Supplementary Fig. S11). Subsequently, the targeted chromosome may evolve to reduce the copy number of the satellite DNA, or to change the satellite DNA sequence that can be transcribed by other transcription factors to avoid being targeted (Supplementary Fig. 9). Repeated cycles of changing satellite DNA sequence/amount and transcription factors/chromatin remodelers may eventually result in incompatibility between satellite DNA and their regulators, leading to speciation.

There are natural variations in satellite DNA abundance at a megabase scale ^23^, which may lead to biased inheritance of chromosome variants, leading to meiotic drive if left unremodeled in spermatocytes. Accordingly, satellite DNA transcription may serve as a mechanism to prevent male meiotic drive. Interestingly, the well-established meiotic driver in *D. melanogaster*, *Segregation Distorter* (*SD)*, involves satellite DNA (*Rsp*) and a RanGAP mutation, where the chromosome II containing a higher amount of *Rsp* satellite DNA is specifically killed with the very same cytological phenotype (sperm DNA compaction defects described in this study)^39–41^. It is tempting to speculate that a RanGAP mutation compromises the nuclear import of proteins responsible for the remodeling of *Rsp* satellite DNA, making the chromosome II with a high amount of *Rsp* satellite DNA fail in sperm DNA packaging. In addition, many of *Drosophila* “hybrid incompatibility loci” that render hybrids lethal or sterile contain satellite DNA (*Zhr*) and encode satellite DNA/heterochromatin-binding proteins (*Hmr, Lhr/HP3*, and *OdsH*)^42–46^. Moreover, rapid changes in the binding specificity of heterochromatin proteins have been documented^46^. The present study may provide a hint as to how satellite DNA sequence and its binding proteins may engage in an evolutionary arms race.

In summary, the present study provides a model to explain the significance of widespread transcription broadly observed in male germline, where it serves to remodel the chromatin of male germline to make it compatible with histone-to-protamine transition. Such requirement for transcription-mediated chromatin remodeling for sperm DNA packaging may provide a potential link between rapid evolution of satellite DNA and male meiotic drive.

## Methods

### Fly Husbandry

All experimental flies were raised on modified Bloomington *Drosophila* Stock Center (BDSC) cornmeal food (agar 0.65%, cornmeal 6.71%, inactivated yeast 1.59%, soy flour 0.92%, corn syrup 7.0%, tegosept 0.15%; without propionic acid; less dry time) at 25°C, and young flies (1-to 3-day-old adults) were used for all experiments. All fly stocks were maintained at room temperature while not in use for experiments, and stocks that exhibited spontaneous spermatid DNA compaction errors were replaced with new stocks. Control flies were either the sibling from the same genetic cross or the parental *GAL4* stock. The following fly stocks were used: *y^1^w^1^* (BDSC 1495), *nos-GAL4:VP16*^47^, *bam-GAL4:VP16*^48^, *Ubi-GFPS65C:α1-tub84B*^49^, *H3.3A-Dendra2* ^50^*, C(1)RM/C(1;Y)6, y^1^w^1^f^1^/O* (BDSC 9460; males from this stock were crossed with *y^1^w^1^* females to generate X/O males), *UAS-TRiP.HMS01699* (BDSC 38255; *UAS-HP2^RNAi^*), *UAS-AAGAG^shRNA^* (gift from Gary Karpen^3^), *Su(var)2-HP2-GFP.FPTB* (BDSC 68184; *hp2-gfp*), *UAS-HP2-S* (this study), *UAS-HP2-L* (this study), Ithaca I16 and Beijing B52 of Global Diversity lines (gift from Andrew Clark^36^). Combinations of transgenes were generated using the CyO; TM6B double balancer strain.

### Immunofluorescence (IF) Staining

Testes were dissected in 1X PBS, and fixed for 30 minutes at room temperature (RT) in 1mL fixative (1X PBS (Invitrogen AM9624; 4mL of 10X PBS), 4% formaldehyde (ThermoFisher Scientific 28908; 10mL of 16% solution), 0.1% Triton X-100 (Millipore Sigma T9284; 400µL of 10% solution) in H_2_O (25.6mL); stored at −20°C). Testes were rinsed with 1mL PBST (1X PBS (Invitrogen AM9624; 5mL of 10X PBS), 0.1% Triton X-100 (Millipore Sigma T9284; 500µL of 10% solution) in H_2_O (44.5mL); stored at 4°C), permeabilized for at least 3 hours at RT in 1mL PBST, blocked for 1 hour at RT in 200µL blocking solution (1X PBS (Invitrogen AM9624; 5mL of 10X PBS), 3% BSA (Fisher Scientific BP1605-100; 1.5g in 25mL H_2_O), 0.1% Triton X-100 (Millipore Sigma T9284; 500µL of 10% solution) in H_2_O (up to 50mL); stored at −20°C), and incubated overnight at 4°C in 200µL blocking solution with primary antibodies. Samples were rinsed with 1mL PBST, washed for at least 1 hour at RT in 1mL PBST, incubated overnight at 4°C in 200µL blocking solution with secondary antibodies, rinsed and washed as above. Samples were mounted in VECTASHIELD with DAPI (Vector Laboratories H-1200). Images were acquired using Leica Stellaris8 confocal microscope with a 63X oil immersion objective lens (NA = 1.4) and processed/analyzed using ImageJ software. The primary antibodies used were anti-Mst77F (1:1000 dilution; guinea pig^51^) and anti-H3K9me3 (1:200 dilution; rabbit; Abcam ab8898). The secondary antibodies used were anti-rabbit IgG conjugated with Alexa Fluor 647 (1:200 dilution; goat; Abcam ab150079) and anti-guinea pig IgG conjugated with Alexa Fluor 568 (1:200 dilution; goat; Abcam ab175714).

### RNA Fluorescent in situ Hybridization (FISH)

Testes were dissected in 1X PBS, and fixed for 30 minutes at RT in 1mL fixative. Testes were washed briefly (5 min x 2) at RT in 1mL PBST, rinsed briefly (5 min) at RT with 1mL wash buffer (2X SSC (Invitrogen AM9770; 5mL of 20X SSC), 10% formamide (Millipore Sigma S4117; 5mL), 0.1% Triton X-100 (Millipore Sigma T9284; 500µL of 10% solution) in H_2_O (39.5mL); stored at 4°C), and hybridized overnight at 37°C in 200µL hybridization buffer (10% dextran sulfate (Millipore Sigma D8906; 1g in 5mL H_2_O), 1mg/mL yeast tRNA (Millipore Sigma R8759; 1mL of 10mg/mL solution), 0.5% BSA (Invitrogen AM2616; 1mL), 2X SSC (Invitrogen AM9770; 1mL of 20X SSC), 10% formamide (Millipore Sigma S4117; 1mL), 2mM Ribonucleoside Vanadyl Complex (NEB S1402; 100µL) in H_2_O (up to 10mL); stored at −20°C). Following hybridization, samples were washed two times in wash buffer for 30 minutes each at 37°C and mounted in VECTASHIELD with DAPI (Vector Laboratories H-1200). Images were acquired using Leica Stellaris8 confocal microscope with a 63X oil immersion objective lens (NA = 1.4) and processed/analyzed using ImageJ software. RNA FISH signal intensity is semi-quantitative due to inherent technical (e.g., different tubes) and biological (e.g., testis thickness) variations. RNA FISH intensities among different genotypes (Figure 2C) were analyzed from images with roughly the same intensity of internal control (GFP-tubulin). Fluorescently labeled probes were added to the hybridization buffer to a final concentration of 50nM (1µL of 10µM probe solution in 200µL hybridization buffer). Probes against the repetitive DNA transcripts were from Integrated DNA Technologies: Cy5-(AAGAG)x6, Cy3-(CTCTT)x6; Cy3-(AATAACATAG)x3, Cy5-(CTATGTTATT)x3; Cy5-(AATAT)x6, Cy3-(ATATT)x6; Cy3-(AACAC)x6, Cy5-(GTGTT)x6; Cy3-(AAGAC)x6, Cy5-(GTCTT)x6. *ORY* probes Quasar 670-(TTTTTGGCTTTCTTTCTGTC, AAAAGTTGAGGCTCCGAGTT, CGTTTAATTCGCGATGCTTC, CTTCATCTACATACCGACGA, TCCGATGTTGAAGTCAGTTC, CCCCCAACAAATCTTTAAGT, TGATGCATCTGATTCTTTCC, TCTTGCACTAACTGTTCTCG, CAGTTGTACCACTATTTCGG, TTAGCTCGAGAAGCCATTTT, ATTCAACGTTTAGGCGTTCG, CCATTGTCTTCATCAAAGCT, TTTCCATGTGCTTCTTTTTG, CTCTATTCGCAATATCCAGT, AGTTTACGTGTCGTTTCTGA, GCATTGCCTTATTTAATGCG, GCCTTAACTGCTTTATCATT, AAGCCTGCTCGTTAATTGAG, TTGTACATTCTTGTTGTCGC, CATCCTTCTCCAGAGAATTA, GACGAATATCATCCGTTCGG, CACGTTGCAATGTCTCTTTG, ATGTTTTAAGTCCGCCATAG, TCTTTCTCTGCTTTGGATTT, CGTCTTCAGAGAGATTTGGT, TCGTGTAACTTTTCCGTGAG, ACGGACTCCAACTGTTTTTT, CGCTGAAGCATCATTTTCTC, TTTTACTTCGTTTGCGCTTA, GCCACTAAGTTTTCTTTGTT, TTTCTCTCATATCCTTACGT, CAATCCGACTAGGTTACGTT, AACAATCTCGTCATTCCGTC, GGGCATTTTGTGCAATTTGA, AAGGCATTCACTTTCAGTGT, CTCATGTCAGCAGTACTTTT, TCTTCAGTTAGAGCTCGTAT, CAACGATGTACTCCTGTAGG, AATTTCTTAGGATCTTCGCC, TATTGACTGCTTTAGGGAGC, AGCCTCACACAGTTTATTTT, GCATATGAGATAGCATCCTT, TAATGAACGCTCTGCTGCTA, GTCTTGATTTAAGTTCCACC, AATTGACAGCTCTCCGATTT, TATTTCGTTTTTCCCATCTG, ATCCACATTCCATATACTCT, ATATCCGTTAACTTCGCACA)

### Quantification of RNA FISH Signal

To quantify RNA FISH signal (for Fig. 1a, 2b, S2f and S3a), we chose spermatocytes of the indicated stages and quantified their signal intensity. To minimize artifactual variations in the signal intensity (such as the depth of the cells within the tissue), images were acquired from near the surface of the tissue, which showed the brightest signal. Z-stacks (step size: 0.4 µm) were collected to cover the entire depth of the spermatocyte to be quantified. To quantify the signals within each spermatocyte, three circular regions per nucleus of 1.2µm in diameter were chosen, while avoiding the areas that contain the signal from the other cells overlapping with the cell of interest. Then, the mean intensity of these regions was calculated after subtracting the cytoplasmic background. These values were then normalized to the mean value of the control.

### Analysis of HP2 ChIP seq for satellite DNA binding

HP2 ChIP-seq data were obtained from the SRA (accessions: SRR870205, SRR870204, SRR870206, SRR870207) (modENCODE ID: 5593)^28^. We analyzed all samples using the k-seek program, and the resulting data were compiled into a summary table using the k.compile.pl script ^36^. Replicate counts were merged for both ChIP-seq and input samples. Satellites with a repeat unit length shorter than 4 bp were excluded. We then selected five major satellites of interest (AAGAG, AACATAGAAT, AATAT, AACAC, AAGAC), while all remaining satellites were combined into a category labeled “other satellites”. For each satellite, enrichment was calculated as the ratio of the satellite’s proportion in the ChIP-seq sample to its proportion in the input. This ratio served as the enrichment metric for each satellite in HP2. The satellite proportion in the input was used as a measure of satellite abundance.

### Reverse Transcribed Quantitative Polymerase Chain Reaction (RT-qPCR)

Dissected testes from 50 flies were pooled in ice-cold 1mL 1X PBS. Samples were first homogenized in ice-cold 300µL TRIzol (Invitrogen 15596026), supplemented with 700µL TRIzol, vortexed vigorously, centrifuged for 5 min at 4°C. Approximately 1mL of clear supernatant was transferred to a new tube, and 400µL of phenol:chloroform (Invitrogen AM9722) was added. Samples were mixed gently by hand for 15 sec, incubated for 3 min at RT, centrifuged for 15 min at 4°C. Approximately 400µL of the top aqueous phase was transferred to a new tube, and 1µL of GlycoBlue Coprecipitant (Invitrogen AM9515), 1mL of 100% ethanol, and 140µL of 5M NH_4_Oac were added. RNA samples were stored at −80°C until the RT reaction. RNA samples were centrifuged for 20 min at 4°C (discard supernatant), washed with 700µL 70% ethanol, centrifuged for 10 min at 4°C (discard supernatant), dried for 15 min at RT, and dissolved in 20µL of 95°C H_2_O. RNA was denatured for 5 min at 65°C and put on ice at least for 1 min. Almost all RNA (20µL, typically 5µg RNA yield) was used for synthesis of cDNA (mixed with 25µL buffer and 5µL SuperScript III (Invitrogen 11752); 25°C 10min, 50°C 30min, 85°C 5min, 4°C keep; mixed with 2.5µL RNase; 37°C 20min; stored at −20°C). Quantitative PCR was done using PowerUp SYBR Green Master Mix (Applied Biosystems A25742; 10µL reaction volume with 5µL of 2X PowerUp SYBR Green Master Mix, 3.5µL H_2_O, 0.5µL cDNA, and 1µL of 5µM forward and reverse primer mix; Primers from IDT were dissolved with H_2_O to make 100µM stock, and 10µL of 100µM forward primer and 10µL of 100µM reverse primer were mixed with 180µL H_2_O to make 5µM primer mix) and assessed with QuantStudio 6 Flex system (Applied Biosystems; 96-well, 0.2mL; standard curve; SYBR Green reagents; standard) with the following PCR condition: 50°C 2 min, 95°C 2 min; 40 cycles of 95°C 15s, 55°C 15s, 72°C 60s (PCR Stage); continuous cycles of 95°C 15s, 55°C 15s, 95°C 15s (Melt Curve Stage). Expression values calculated from the 2^−ΔΔCt^ comparative Ct method, 2^−[(Ct^Target, Experimental^ – Ct^Housekeeping, Experimental^) – (Ct^Target, Control^ – Ct^Housekeeping, Control^)]. Primers used for this study are as follows: HP2-S Fwd 5’-ATTTGGGTGACTTATACGGGGACT-3’, HP2-L Fwd 5’-ATCCCGAACCCAGCACAT-3’, HP2 Rev 5’-CTGGTTTAACTTGTTCCTTTTTGCCAGT-3’, Und (as housekeeping) Fwd 5’-GCAAGAAAAGCGGTCAGACT-3’, Und (as housekeeping) Rev 5’-CGTGTTGATACGGTCCAGAG-3’, kl-5 Fwd 5’-TGTCTATGAGTGCCCAGTTTAC-3’, kl-5 Rev 5’-GTCCACTTACTTGCTCTCTCTC-3’, kl-3 Fwd 5’-ATGGTGCTGGTTGGGATAAG-3’, kl-3 Rev 5’-TTGGCAGCCGTTGAGAAT-3’, kl-2 Fwd 5’-TTTACCAGTGTCCCGCATATT-3’, kl-2 Rev 5’-AGTGCAGTACCTCGCTTTATC-3’, ORY (Last Exon) Fwd 5’-CAGCGTATAAACCGACAAATGG-3’, ORY (Last Exon) Rev 5’-GACAGCTCTCCGATTTCACTAA-3’, ORY (Exons 4-5) Fwd 5’-ACTGTGCACTTCCCTTTGT-3’, ORY (Exons 4-5) Rev 5’-GAGATGAAATGGCGCAAGAAAT-3’, CCY Fwd 5’-CGGAGCCGTAAAGGATGATT-3’, CCY Rev 5’-CGCTGACCTGATAACCCTATTC-3’, DIP-λ Fwd 5’-ACACCCTGGATACTCACAAATG-3’, DIP-λ Rev 5’-AGTGAAACACACGCCAGAA-3’, CG44666 Fwd 5’-GCCGATTTGTGCAGTCTTTC-3’, CG44666 Rev 5’-GGACGATGTGGATGCTGTTA-3’, Myo81F Fwd 5’-CCTCTTCATCGAGCGACTATTT-3’, Myo81F Rev 5’-ACGATGTTCCAGTAGGTGAATAC-3’, Pzl Fwd 5’-GGATGATATCCTACTACCAGCTTTG-3’, Pzl Rev 5’-CCGGATAACAAGCCTCCTTTAT-3’, Mitf Fwd 5’-TCAGATGACTGTTGCGGTATAA-3’, Mitf Rev 5’-ATACCCGTGTGGTGAGAATG-3’.

### DNA FISH

Adult testes (for meiotic chromosome spread, typically three flies per slide) or third instar larval brains (for mitotic chromosome spread, typically three larvae per slide) were dissected in PBS. Samples were fixed for 4 min at RT in 50 µL of modified fixative (45% acetic acid (450µL of glacial acetic acid) and 2.5% formaldehyde (550µL of fixative described earlier)) placed on Superfrost Plus microscope slides (ThermoFisher Scientific 22-037-246). Samples were squashed under a coverslip, and then immediately frozen in liquid nitrogen. Coverslips were quickly removed and slides were immediately dehydrated in a Coplin jar filled with 100% ethanol. Then, slides were dried at RT at least for 1 hour. 17 µL of hybridization buffer (10% dextran sulfate (Millipore Sigma D8906; 1g in 2mL H_2_O), 50% formamide (Millipore Sigma S4117; 5mL), 2X SSC (Invitrogen AM9770; 1mL of 20X SSC), 10mM EDTA (Invitrogen AM9260G; 200µL of 0.5M EDTA) in H_2_O (up to 8.5mL); stored at −20°C) was mixed with 3µL probe mix (0.5 µM probe (1µL of 10µM probe solution) up to three colors or substitute with H_2_O). This hybridization mix was applied to samples on a slide and covered with a cover slip. Samples were incubated at 95°C for 5 min, cooled and wrapped in parafilm, then incubated overnight at RT in a dark humid chamber. In a Coplin jar filled with 0.1X SSC (5mL of 20X SSC diluted with H_2_O up to 1L), coverslips were removed and slides were washed three times for 15 min each. Slides were dried, sample locations were marked, and samples were mounted in VECTASHIELD with DAPI (Vector Laboratories H-1200). For the whole testis staining, testes were dissected in 1X PBS, fixed for 30 minutes at RT in 1mL modified fixative (1mM EDTA (Invitrogen AM9260G; 2µL of 0.5M EDTA) and 1mL of fixative described earlier), rinsed with 1mL PBST-EDTA (1X PBS (Invitrogen AM9624; 5mL of 10X PBS), 0.1% Triton X-100 (Millipore Sigma T9284; 500µL of 10% solution), 1mM EDTA (Invitrogen AM9260G; 100µL of 0.5M EDTA) in H_2_O (44.4mL); stored at RT), washed for 1hr at RT in 1mL PBST-EDTA, rinsed with 1mL PBST (no EDTA), incubated for 10 min at 37°C in 100µL RNase A (2mg/mL in PBST), washed for 10 min at RT in 1mL PBST-EDTA, rinsed with 1mL SCCT-EDTA (50mL SCCT (5mL of 20X SSC and 250µL of 20% Tween in 45mL H_2_O; stored at RT) and 100µL of 0.5M EDTA; stored at RT), washed for 15 min at RT in 100µL SCCT-20%F (800µL SCCT and 200µL formamide, freshly made), washed for 15 min at RT in 100µL SCCT-40%F (600µL SCCT and 400µL formamide, freshly made), washed for 15 min at RT in 100µL SCCT-50%F (500µL SCCT and 500µL formamide, freshly made), washed for 30 min at RT in 100µL SCCT-50%F, and put in 100µL hybridization mix (85µL hybridization buffer mixed with 5µL of 10µM probe up to three colors or substitute with H_2_O). Samples were denatured for 5 min at 95°C and hybridized overnight at 37°C. Following hybridization, samples were washed two times in 1mL SCCT-EDTA for 30 minutes each at RT and mounted in VECTASHIELD with DAPI (Vector Laboratories H-1200). Images were acquired using Leica Stellaris8 confocal microscope with a 63X oil immersion objective lens (NA = 1.4) and processed/analyzed using ImageJ software. Probes used for this study are as follows: Cy3-(TAGA)x8, Cy3/Cy5-(AATAC)x6, Cy5/AlexaFluor-488-(AAGAG)x6.

### Genotyping

X/Y; II+/II^ΔAAGAG^ (*bam>HP2^RNAi^*) were mated with *y^1^w^1^* females for 5 days, and the eclosed flies was collected a week later for genotyping. The *y^+^* transgene on II^+^ was used to genotype male offspring by the *y^+^* body color. Because X/Y; II^+^/II^ΔAAGAG^ (*bam>HP2^RNAi^*) carry a wild-type X chromosome with the endogenous *y^+^* gene, female offspring was genotyped by PCR using *y^+^* transgene-specific sequences. Each fly was homogenized in 50µL extraction buffer (1X TE (Sigma Aldrich 93283; 49µL), 25mM NaCl (0.5µL of 2.5M NaCl), 200µg/mL proteinase K (0.5µL of 20mg/µL), incubated for 30 min at 37°C followed by 1 min at 95°C. Genomic DNA was stored in −20°C. 1µL of genomic DNA was added in PCR mixture (DreamTaq DNA Polymerase (ThermoFisher Scientific EP0705; 0.2µL), 0.5µM primer (2µL of 5µM forward and reverse primer mix), 200µM dNTP (0.4µL of 10mM dNTP), DreamTaq Green Buffer (ThermoFisher Scientific B71; 2µL) in H_2_O (14.4µL)). Target is amplified by touchdown PCR (95°C 3 min; 15 cycles of 95°C 30s, 70°C (−1°C/cycle) 45s, 72°C 30s; 20 cycles of 95°C 30s, 55°C 45s, 72°C 30s; 72°C 5 min and 10°C keep). 10µL of PCR reaction was analyzed by gel electrophoresis of 1.5% TAE gel 120V for 45min with 2µL of 100bp ladder. Primers used for this study are as follows: Fwd 5’-CTAGAGTAAGTAGTTCGCCAGTTAAT-3’ and Rev 5’-GCTGAATGAAGCCATACCAAAC-3’.

### Fertility Assay and sex ratio scoring

One male (control or *bam>HP2^RNAi^*) and three *y^1^w^1^* virgin females (1-3 days old) are mated for 4 days in the 25°C incubator. After 4 days, parents were discarded and the number of eclosed flies is counted a week later. After 4 days, parents were discarded and the numbers of eclosed male and female progeny was counted for each mating pair. The sex ratio was calculated as the number of female offspring/the total number of offspring.

### Transgenic Fly Generation

The codon-modified fragment of HP2 (RNAi target sequence CAGGAGAGGAATGAAGAACAA was changed to CAAGAACGCAACGAGGAGCAG) was synthesized by the IDT gBlocks Gene Fragments service (1000ng; suspended in 100µL TE to make 10ng/µL solution and incubated at 50°C for 20 minutes; stored at −20°C) and cloned into UFO12514 (pUAST-based vector with HP2-S cDNA, obtained from *Drosophila* Genomics Resource Center (DGRC) Stock 1648698) using In-Fusion cloning. Briefly, the UFO12514 vector was amplified by standard PCR (95°C 2 min; 30 cycles of 95°C 20s, 55°C 20s, 72°C 30s/kb using Herculase II Fusion DNA Polymerase; 72°C 3 min and 10°C keep); 0.5µL vector (at least 50ng), 0.5µL insert (at least a few fold excess of insert), 3µL H_2_O, and 1µL In-Fusion Snap Assembly Master Mix (Takara Bio 638947) were mixed and incubated for 15 min at 50°C followed by 4°C keep; 50µL DH5α (NEB C2987) and 5µL In-Fusion reaction mixture was mixed and incubated for 15 min on ice, and the bacteria was heat shocked for 45 sec at 42°C, on ice for 1 min, recovered in 200µL SOC (briefly if ampicillin resistant, 1h at 37°C if non-ampicillin resistant), and cultured on a LB plate with 100µg/mL carbenicillin or other antibiotics overnight at 37°C; Each colony is dissolved in 10µL H_2_O, and 1µL was used for colony PCR (mixed with19µL DreamTaq PCR mixture used for genotyping followed by touchdown PCR that amplify the insert) and 9µL was added in 200µL LB (stored at 4°C during colony PCR, and cultured in 3mL LB overnight at 37°C); 500µL of cultured *E. coli* was mixed with 500µL of 50% glycerol and stored at −80°C, and the rest of cultured *E. coli* was used to collect plasmids using Miniprep (QIAGEN 27104; culture in 2mL LB overnight; centrifuge for 3 min; dissolve in 250µL RNase solution; add 250µL SDS solution; gently invert 5 times; incubate at RT for 3min; add 350µL neutralizing solution; gently invert 5 times; centrifuge for 10min; transfer 700µL supernatant to the spin column; add 700µL salt wash solution; centrifuge for 30sec; add 700µL ethanol wash solution; centrifuged for 30sec; centrifuge for another 1min; elute in 50µL water); The entire plasmid was sequenced by Plasmidsaurus Inc; Small cloning errors were corrected by QuikChange Lightning Site-Directed Mutagenesis Kit (Agilent 210518). HP2-L fragments were amplified from testis cDNA library: Briefly, 1µL of cDNA and 5µL of 5µM forward and reverse primer mix for each fragment were added in PCR mixture (Herculase II Fusion DNA Polymerase (Agilent 600675; 1µL), 250µM dNTP (1.25µL of 10mM dNTP), 5x Buffer (10µL) in H_2_O (31.75µL)). For each fragment, two tubes with 50µL PCR reaction were prepared and amplified by touchdown PCR, adjusting the extension time (30s/kb). The PCR products were run on 0.7% TAE gel and extracted by gel purification (QIAGEN 28506; mix gel and binding solution in 1:1 ratio; 50°C for 5-10min; spin down; add 700µL binding solution; centrifuge for 1min; add 700µL ethanol wash solution; centrifuge for 1min; centrifuge for another 1min; elute in 20µL water). The two HP2-L fragments were fused by overlap extension PCR: Briefly, 20µL PCR reaction (0.5µL of Herculase II Fusion DNA Polymerase, 0.5µL of 10mM dNTP, 4µL of 5x Buffer, ([length of fragment (bp) ÷ 10] ng of each fragment with H_2_O) was incubated in a thermocycler (95°C 3 min; 14 cycles of 95°C 30s, 72°C (−0.5°C/cycle) 45s, 72°C 30s/kb; 72°C 5 min and 10°C keep). 30µL spike-in mixture (17.75µL H_2_O, 5µL of 5µM forward and reverse primer mix for the entire fragment, 6µL of 5x Buffer, 0.75µL of 10mM dNTP, 0.5µL of Herculase II Fusion DNA Polymerase) was added in each PCR reaction (total 50µL) and incubated in a thermocycler (95°C 3 min; 14 cycles of 95°C 30s, 72°C (−0.5°C/cycle) 45s, 72°C 30s/kb; 20 cycles of 95°C 30s, 65°C 45s, 72°C 30s/kb; 72°C 5 min and 10°C keep). The fused product was gel purified in 20µL H_2_O and inserted into pCR-Blunt II-TOPO vector (ThermoFisher Scientific 450245; 4µL gel purified DNA, 1µL salt solution, and 1µL TOPO vector were incubated for 5 min at RT; 5µL reaction was transformed in 50µL DH5α and cultured on a LB plate with 50µg/mL kanamycin). The fused HP2-L fragment was subcloned into the HP2-S plasmid (amplified by long template (>10kb) PCR: 50µL Herculase PCR mixture was incubated in a thermocycler (92°C 2 min; 10 cycles of 92°C 20s, 55°C 20s, 68°C 30s/kb; 20 cycles of 92°C 20s, 55°C 20s, 68°C 30s/kb +20s/cycle; 68°C 8 min and 10°C keep)) using In-Fusion cloning. The PhiC31 integrase-mediated integration was conducted by BestGene Inc. The pUAST construct with RNAi resistant HP2-S is inserted into the VK27 site on 3R (BDSC 9744). The pUAST construct with RNAi resistant HP2-L is inserted into the attP2 site on 3L (BDSC 8622). Both constructs were balanced with TM3.

### Analysis of spermatocyte-enriched DNA-binding proteins

The list of proteins annotated as transcription factors or as containing a DNA-binding domain were obtained from FlyBase, using the associated “Gene Group” lists. These lists were merged and redundancies removed. The Fly Cell Atlas testis dataset^1^ was used to identify spermatocyte-enriched genes using the FindMarkers tool in Seurat v5.1 with default paremeters, with spermatocytes defined as per the previous annotation (clusters C-G, as in Figure 4A, Raz et al^1^). Bona fide markers were considered those with an adjusted p-value < 10^-9^. These lists were then intersected to obtain a catalog of putative transcription factors in spermatocytes.

### Comparison of transcribed and translated protein-coding genes

Mass-spec data from Gärtner 2019 (accessed from Project PXD010627 Proteomics Identifications Database, folder MaxQuant_Output.zip, subfolder proteinGroups.txt) was sorted by annotated “protein groups”. We binned the data with any protein group with a specific corresponding gene ID and at least one peptide across in all three adult testis replicates considered “translated”, and all others considered absent. Protein presence in non-adult stages was not considered. Genes were considered transcribed if they had an average expression of >=0.1 ln(UMIs-per-10,000 + 1) across the entire testis sequencing dataset^1^. These lists were intersected to find genes transcribed (present in the sequencing dataset but not the mass-spec dataset), translated (present in the mass-spec dataset but not the sequencing dataset), neither or both.

### Identification of transcribed transposable elements and lncRNAs

We generated a new reference file that contains both the DM6 reference and the consensus sequences of *Drosophila* transposable elements (https://github.com/bergmanlab/drosophila-transposons). Specifically, a .gtf file was generated from the consensus TE .fa file (D_mel_tranasposon_sequence_set.fa) with every TE sequence considered an “exon”. New .fa and .gtf files were then generated by concatenating the respective files from DM6 and the TE consensus. The Fly Cell Atlas testis dataset was then remapped to this new reference using default parameters in the cellranger 8.0.1 tool. The resulting mapped file was processed, dimensionality reduced, and clustered according to previous parameters^1^; and annotations were transferred from the previous publication. lncRNAs, as identified in DM6, or TEs, as identified in the consensus file, were considered “expressed” in spermatocytes if they had an average expression of >=15 ln(UMIs-per-10,000 + 1) across all spermatocytes, with spermatocytes annotated as above.

### Statistics and Reproducibility

Information of statistics is provided in respective figure legends or Source Data file. For some imaging results where quantification is not provided, experiments were repeated at least 2 times, each with 10 pairs of testes, and each testis contained hundreds of spermatocytes (Figure 1i, 3e).

#### Data availability statement

The previously published data sets utilized are, Raz et al^1^, NCBI Gene Expression Omnibus GSE220615; Gartner et al.^21^, PXD010627 (ProteomeCentral), and modEncode (2010)^28^ (www.modencode.org/publications/integrative_fly_2010/) as detailed in ^28^. Source Data are available with this paper.

## Acknowledgements

We thank Dr. Gary Karpen, and Dr. Andrew Clark for reagents, and the members of the Yamashita lab for discussions and comments on the manuscript. We thank the FlyBase, Bloomington *Drosophila* Stock Center, and *Drosophila* Genomics Resource Center for reagents and information.

## Funding

This research is funded by the Howard Hughes Medical Institute, Gordon and Betty Moore Foundation, Whitehead Institute for Biomedical Research (to Y.M.Y), and National Institute of General Medical Sciences (F32GM143850 and K99GM154143 to A.A.R).

## Author Contributions

Conceptualization, T.K. and Y.M.Y.; methodology, T.K.; investigation, T.K. and M.N-D; software, A.R. and R. L.; resources, J.M.F. and Y.M.Y.; writing – original draft, T.K.; writing – editing, T.K. and Y.M.Y.; funding acquisition, Y.M.Y.; supervision, Y.M.Y.

## Competing Interests

The authors declare no competing interests.

## Notes

### Competing Interest Statement

The authors have declared no competing interest.

### Summary of Updates

Additional strain of delta-AAGAG was added to the experiments. Image quantification was added for some images. Better representative images were added.

## References

1. Raz, A.A. et al. Emergent dynamics of adult stem cell lineages from single nucleus and single cell RNA-Seq of Drosophila testes. Elife 12 (2023).

2. Wen, K. et al. Critical roles of long noncoding RNAs in Drosophila spermatogenesis. Genome Res 26, 1233–1244 (2016).

3. Mills, W.K., Lee, Y.C.G., Kochendoerfer, A.M., Dunleavy, E.M. & Karpen, G.H. RNA from a simple-tandem repeat is required for sperm maturation and male fertility in Drosophila melanogaster. Elife 8 (2019).

4. Lawlor, M.A., Cao, W. & Ellison, C.E. A transposon expression burst accompanies the activation of Y-chromosome fertility genes during Drosophila spermatogenesis. Nat Commun 12, 6854 (2021).

5. Soumillon, M. et al. Cellular source and mechanisms of high transcriptome complexity in the mammalian testis. Cell Rep 3, 2179–2190 (2013).

6. Chen, Y. et al. Single-cell RNA-seq uncovers dynamic processes and critical regulators in mouse spermatogenesis. Cell Res 28, 879–896 (2018).

7. Guo, J. et al. The adult human testis transcriptional cell atlas. Cell Res 28, 1141–1157 (2018).

8. Wang, D. et al. A deep proteome and transcriptome abundance atlas of 29 healthy human tissues. Mol Syst Biol 15, e8503 (2019).

9. Xia, B. et al. Widespread Transcriptional Scanning in the Testis Modulates Gene Evolution Rates. Cell 180, 248–262 e221 (2020).

10. Liu, H. & Zhang, J. Higher Germline Mutagenesis of Genes with Stronger Testis Expressions Refutes the Transcriptional Scanning Hypothesis. Molecular biology and evolution 37, 3225–3231 (2020).

11. Walker, P.M. Origin of satellite DNA. Nature 229, 306–308 (1971).

12. Yunis, J.J. & Yasmineh, W.G. Heterochromatin, satellite DNA, and cell function. Structural DNA of eucaryotes may support and protect genes and aid in speciation. Science 174, 1200–1209 (1971).

13. Lohe, A.R., Hilliker, A.J. & Roberts, P.A. Mapping simple repeated DNA sequences in heterochromatin of Drosophila melanogaster. Genetics 134, 1149–1174 (1993).

14. Shaffer, C.D. et al. Heterochromatin protein 2 (HP2), a partner of HP1 in Drosophila heterochromatin. Proc Natl Acad Sci U S A 99, 14332–14337 (2002).

15. Shaffer, C.D. et al. The large isoform of Drosophila melanogaster heterochromatin protein 2 plays a critical role in gene silencing and chromosome structure. Genetics 174, 1189–1204 (2006).

16. Stephens, G.E., Xiao, H., Lankenau, D.H., Wu, C. & Elgin, S.C. Heterochromatin protein 2 interacts with Nap-1 and NURF: a link between heterochromatin-induced gene silencing and the chromatin remodeling machinery in Drosophila. Biochemistry 45, 14990–14999 (2006).

17. Stephens, G.E., Slawson, E.E., Craig, C.A. & Elgin, S.C. Interaction of heterochromatin protein 2 with HP1 defines a novel HP1-binding domain. Biochemistry 44, 13394–13403 (2005).

18. Rathke, C., Baarends, W.M., Awe, S. & Renkawitz-Pohl, R. Chromatin dynamics during spermiogenesis. Biochim Biophys Acta 1839, 155–168 (2014).

19. Loppin, B. & Berger, F. Histone Variants: The Nexus of Developmental Decisions and Epigenetic Memory. Annu Rev Genet 54, 121–149 (2020).

20. Li, H. et al. Fly Cell Atlas: A single-nucleus transcriptomic atlas of the adult fruit fly. Science 375, eabk2432 (2022).

21. Gartner, S.M.K. et al. Stage-specific testes proteomics of Drosophila melanogaster identifies essential proteins for male fertility. Eur J Cell Biol 98, 103–115 (2019).

22. Jagannathan, M., Warsinger-Pepe, N., Watase, G.J. & Yamashita, Y.M. Comparative Analysis of Satellite DNA in the Drosophila melanogaster Species Complex. G3 (Bethesda) 7, 693–704 (2017).

23. Wei, K.H. et al. Variable Rates of Simple Satellite Gains across the Drosophila Phylogeny. Molecular biology and evolution 35, 925–941 (2018).

24. Chang, C.H. & Larracuente, A.M. Heterochromatin-Enriched Assemblies Reveal the Sequence and Organization of the Drosophila melanogaster Y Chromosome. Genetics 211, 333–348 (2019).

25. Bonaccorsi, S., Pisano, C., Puoti, F. & Gatti, M. Y chromosome loops in Drosophila melanogaster. Genetics 120, 1015–1034 (1988).

26. Bonaccorsi, S. & Lohe, A. Fine mapping of satellite DNA sequences along the Y chromosome of Drosophila melanogaster: relationships between satellite sequences and fertility factors. Genetics 129, 177–189 (1991).

27. Fingerhut, J.M., Moran, J.V. & Yamashita, Y.M. Satellite DNA-containing gigantic introns in a unique gene expression program during Drosophila spermatogenesis. PLoS genetics 15, e1008028 (2019).

28. mod, E.C., et al. Identification of functional elements and regulatory circuits by Drosophila modENCODE. Science 330, 1787–1797 (2010).

29. Lu, B.Y., Emtage, P.C., Duyf, B.J., Hilliker, A.J. & Eissenberg, J.C. Heterochromatin protein 1 is required for the normal expression of two heterochromatin genes in Drosophila. Genetics 155, 699–708 (2000).

30. Wei, X., Eickbush, D.G., Speece, I. & Larracuente, A.M. Heterochromatin-dependent transcription of satellite DNAs in the Drosophila melanogaster female germline. Elife 10 (2021).

31. Castel, S.E. & Martienssen, R.A. RNA interference in the nucleus: roles for small RNAs in transcription, epigenetics and beyond. Nat Rev Genet 14, 100–112 (2013).

32. Hoskins, R.A. et al. The Release 6 reference sequence of the Drosophila melanogaster genome. Genome Res 25, 445–458 (2015).

33. Carvalho, A.B., Lazzaro, B.P. & Clark, A.G. Y chromosomal fertility factors kl-2 and kl-3 of Drosophila melanogaster encode dynein heavy chain polypeptides. Proceedings of the National Academy of Sciences of the United States of America 97, 13239–13244 (2000).

34. Hafezi, Y., Sruba, S.R., Tarrash, S.R., Wolfner, M.F. & Clark, A.G. Dissecting Fertility Functions of &lt;em&gt;Drosophil&lt;/em&gt;a &lt;em&gt;Y&lt;/em&gt; Chromosome Genes with CRISPR. Genetics 214, 977 (2020).

35. Carvalho, A.B., Dobo, B.A., Vibranovski, M.D. & Clark, A.G. Identification of five new genes on the Y chromosome of Drosophila melanogaster. Proceedings of the National Academy of Sciences of the United States of America 98, 13225–13230 (2001).

36. Wei, K.H., Grenier, J.K., Barbash, D.A. & Clark, A.G. Correlated variation and population differentiation in satellite DNA abundance among lines of Drosophila melanogaster. Proc Natl Acad Sci U S A 111, 18793–18798 (2014).

37. Ridges, J.T., et al. Overdrive is essential for targeted sperm elimination by Segregation Distorter. bioRxiv (2024).

38. Courret, C., Wei, X. & Larracuente, A.M. New perspectives on the causes and consequences of male meiotic drive. Curr Opin Genet Dev 83, 102111 (2023).

39. Merrill, C., Bayraktaroglu, L., Kusano, A. & Ganetzky, B. Truncated RanGAP encoded by the Segregation Distorter locus of Drosophila. Science 283, 1742–1745 (1999).

40. Larracuente, A.M. & Presgraves, D.C. The selfish Segregation Distorter gene complex of Drosophila melanogaster. Genetics 192, 33–53 (2012).

41. Herbette, M. et al. Distinct spermiogenic phenotypes underlie sperm elimination in the Segregation Distorter meiotic drive system. PLoS genetics 17, e1009662 (2021).

42. Maheshwari, S. & Barbash, D.A. The genetics of hybrid incompatibilities. Annu Rev Genet 45, 331–355 (2011).

43. Presgraves, D.C. & Meiklejohn, C.D. Hybrid Sterility, Genetic Conflict and Complex Speciation: Lessons From the Drosophila simulans Clade Species. Front Genet 12, 669045 (2021).

44. Ferree, P.M. & Barbash, D.A. Species-specific heterochromatin prevents mitotic chromosome segregation to cause hybrid lethality in Drosophila. PLoS Biol 7, e1000234 (2009).

45. Satyaki, P.R., et al. The hmr and lhr hybrid incompatibility genes suppress a broad range of heterochromatic repeats. PLoS genetics 10, e1004240 (2014).

46. Bayes, J.J. & Malik, H.S. Altered heterochromatin binding by a hybrid sterility protein in Drosophila sibling species. Science 326, 1538–1541 (2009).

47. Van Doren, M., Williamson, A.L. & Lehmann, R. Regulation of zygotic gene expression in Drosophila primordial germ cells. Curr Biol 8, 243–246 (1998).

48. Chen, D. & McKearin, D.M. A discrete transcriptional silencer in the bam gene determines asymmetric division of the Drosophila germline stem cell. Development 130, 1159–1170 (2003).

49. Rebollo, E., Llamazares, S., Reina, J. & Gonzalez, C. Contribution of Noncentrosomal Microtubules to Spindle Assembly in Drosophila Spermatocytes. PLoS Biol 2, E8 (2004).

50. Shindo, Y. & Amodeo, A.A. Dynamics of Free and Chromatin-Bound Histone H3 during Early Embryogenesis. Curr Biol 29, 359–366 e354 (2019).

51. Park, J.I., Bell, G.W. & Yamashita, Y.M. Derepression of Y-linked multicopy protamine-like genes interferes with sperm nuclear compaction in D. melanogaster. Proc Natl Acad Sci U S A 120, e2220576120 (2023).

52. Misra, S. et al. Annotation of the Drosophila melanogaster euchromatic genome: a systematic review. Genome Biol 3, RESEARCH0083 (2002).

53. Merel, V., Boulesteix, M., Fablet, M. & Vieira, C. Transposable elements in Drosophila. Mobile DNA 11, 23 (2020).

