## supplementary figures for "Defective transcription of AAGAG satellite DNA causes sex-ratio meiotic drive in *Drosophila*"

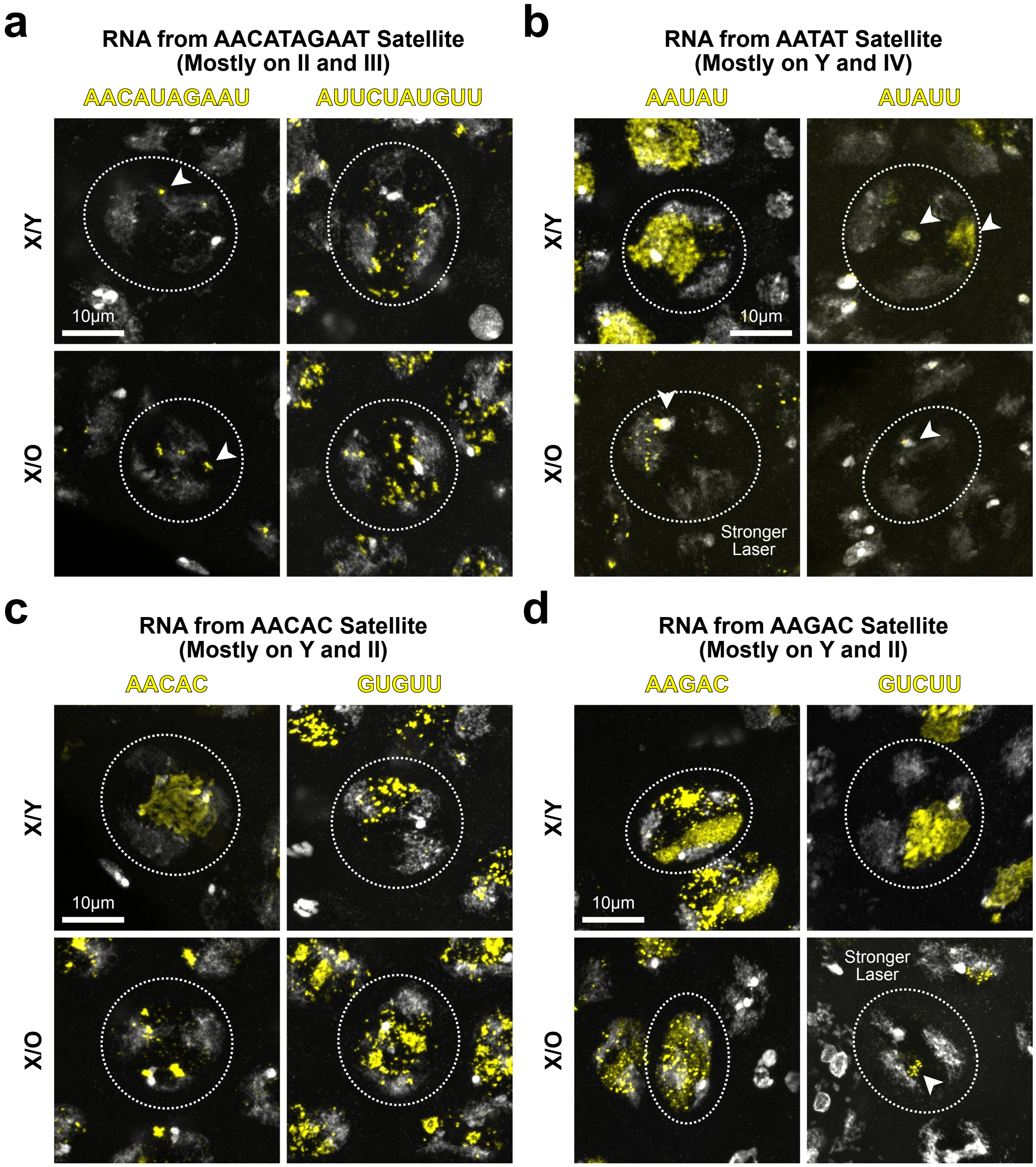


**Supplementary Fig. S1: A broad range of satellite DNAs is transcribed in spermatocytes**

(a-d) RNA FISH of various satellite transcripts in X/Y and X/O spermatocytes. Weakly transcribed loci are highlighted with white arrowheads. Transcripts of indicated satellite DNA are shown in yellow, with DAPI in gray. Experiments were duplicated, each containing >10 pairs of testes.

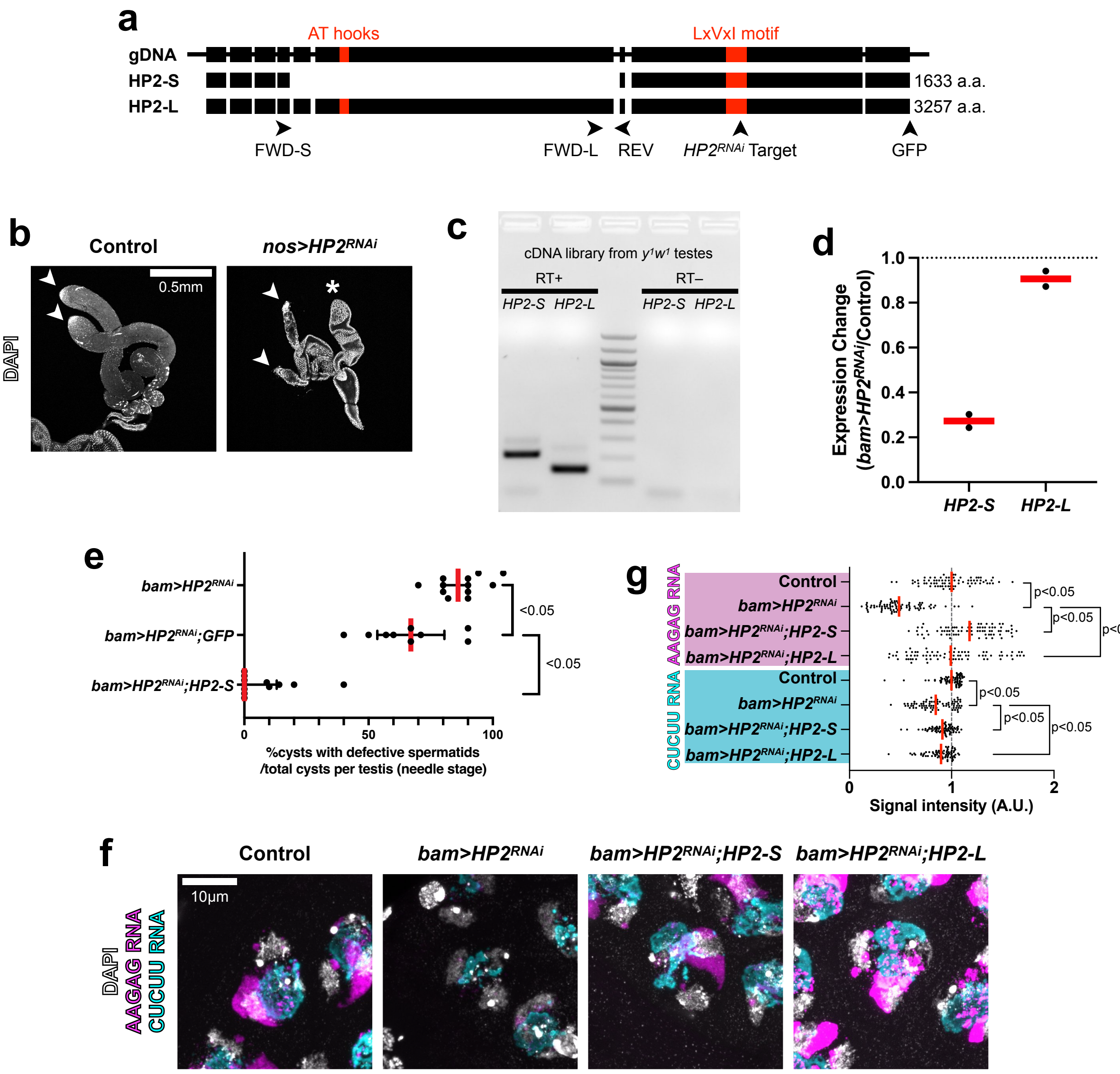


**Supplementary Fig. S2: Both HP2 isoforms can rescue the HP2 RNAi phenotype**

(a) Schematics of the HP2 genomic sequence and the two isoforms (HP2-S and HP2-L). Black boxes are exons, ordered from 5' (left) to 3' (right), with introns, 5' upstream and 3' downstream sequences represented as a line. Satellite DNA interacting domain (two AT hooks, GRP) and HP1-interacting domain (LxVxI motif, LCVKI) are shown in red. Locations of primers used for RT-qPCR (panels c, d), position of GFP-tag, and *HP2*$^{RNAi}$ target sequence (CAGGAGAGGAATGAAGAACAA) are shown with arrowheads.

(b) The testes from control vs. *nos>HP2*$^{RNAi}$ (arrowheads) stained with DAPI. Asterisks show accessory glands. *nos>HP2*$^{RNAi}$ led to the loss of germ cells and degenerated testis, consistent with known HP2's requirement in the viability of mitotically proliferating cell. In this study, *bam>HP2*$^{RNAi}$ was used to study the function of HP2 in spermatocytes.

(c) RT-PCR to detect long and short isoforms of HP2 in the testis.

(d) RT-qPCR using the primer sets for each HP2 isoform in *bam>HP2*$^{RNAi}$ compared to control with technical duplicates. *bam>HP2*$^{RNAi}$ reduced HP2-S expression, but not HP2-L, suggesting that HP2-S is the dominant isoform in spermatocytes. HP2-L is likely expressed in early germ cells (spermatogonia), thus not effectively depleted by *bam>HP2*$^{RNAi}$.

(e) HP2-S transgene (RNAi resistant) rescues spermatid DNA compaction defects in *bam>HP2*$^{RNAi}$. UAS-GFP did not rescue *bam>HP2*$^{RNAi}$ phenotype, demonstrating that the rescue by UAS-HP2-S is not due to dilution of *bam-gal4* driver. (n = 12 testes for *bam>HP2*$^{RNAi}$, n = 9 testes for *bam>HP2*$^{RNAi}$ *UAS-GFP,* n = 12 for *bam>HP2*$^{RNAi}$ *UAS-HP2-S*). Each dot represents an individual testis; red line, median; black lines, interquartile range. The p-value was calculated by the Mann-Whitney U test (two-tailed): *bam>HP2*$^{RNAi}$ vs *bam>HP2*$^{RNAi}$ *UAS-GFP*, p = 0.0078; *bam>HP2*$^{RNAi}$ *UAS-GFP* vs *bam>HP2*$^{RNAi}$ *UAS-HP2-S*, p < 0.0001, and a p-value of less than 0.05 is judged as statistically significant. Exact numbers of samples can be found in the Source Data file.

(f) RNA FISH of AAGAG/CUCUU satellite transcripts in control, *bam>HP2*$^{RNAi}$, and *bam>HP2*$^{RNAi}$ with HP2 rescue constructs (designed to be resistant to RNAi, see Method). Both HP2-S and HP2-L can rescue the phenotype.

(g) Relative signal intensities of AAGAG/CUCUU transcripts in control, *bam>HP2*$^{RNAi}$, and *bam>HP2*$^{RNAi}$ with HP2 rescue construct spermatocytes (n= 77, 81, 78, 78, 79, 79, 78, 78 spermatocytes from top to bottom genotypes). Red line, mean. Each dot represents the RNA FISH intensity in a single spermatocyte (see Methods). The p-value was calculated by unpaired t test (two-tailed). For AAGAG RNA: control vs *bam>HP2*$^{RNAi}$, p < 0.0001; *bam>HP2*$^{RNAi}$ vs *bam>HP2*$^{RNAi}$ *UAS-HP2-S*, p < 0.0001; *bam>HP2*$^{RNAi}$ vs *bam>HP2*$^{RNAi}$ *UAS-HP2-L*, p < 0.0001. For CUCUU RNA: control vs *bam>HP2*$^{RNAi}$, p < 0.0001; *bam>HP2*$^{RNAi}$ vs *bam>HP2*$^{RNAi}$ *UAS-HP2-S*, p = 0.0045; *bam>HP2*$^{RNAi}$ vs *bam>HP2*$^{RNAi}$ *UAS-HP2-L*, p = 0.0323, and a p-value of less than 0.05 is judged as statistically significant. Exact numbers of samples can be found in the Source Data file.

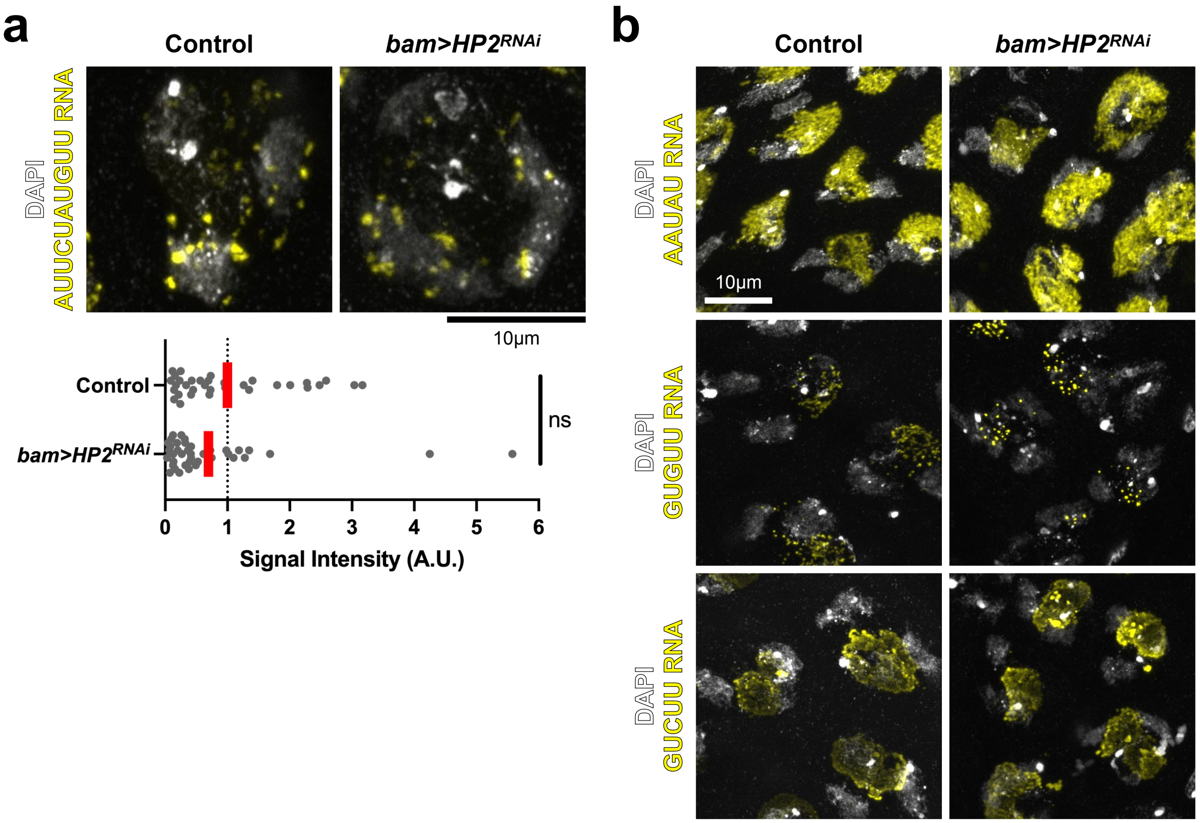


**Supplementary Fig. S3: HP2 depletion has little effect on transcripts from other satellite DNA than AAGAG.**

(a) RNA FISH (top) and relative signal intensities (bottom) of AUUCUACGCC RNA in control and *bam>HP2*$^{RNAi}$ spermatocytes (n = 33 spermatocytes for control, 39 spermatocytes for *bam>HP2*$^{RNAi}$ from ≥14 testes from each genotype). Red line, mean. The p-value was calculated by unpaired t test (two-tailed): p = 0.2083 (ns, not significant). Exact numbers of samples can be found in the Source Data file.

(b) RNA FISH of AAUAU, GUGUU, and GUCUU RNA in control and *bam>HP2*$^{RNAi}$ spermatocytes.

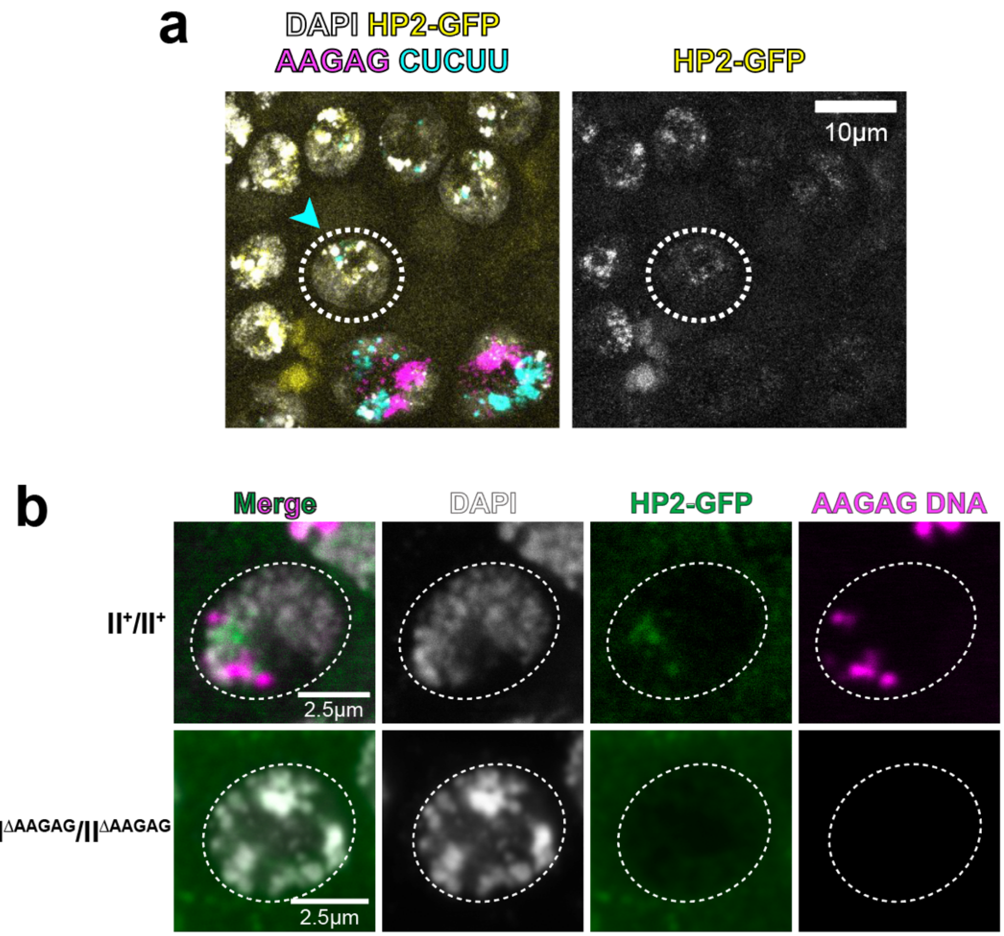


**Supplementary Fig. S4: HP2 is expressed in developing male germ cells and localizes to the chromatin in an AAGAG satellite DNA-dependent manner**

(a) HP2-GFP expression in male germ cells. HP2-GFP was most strongly expressed in spermatogonia (circled, blue arrowhead), prior to strong AAGAG/CUCUU expression in spermatocytes (left bottom cells).

(b) In spermatogonia, HP2-GFP was observed on chromatin near AAGAG satellite DNA. This localization is dependent on AAGAG satellite DNA, as the strain with little AAGAG satellite DNA (Ithaca 16 strain[1]) exhibited an undetectable HP2-GFP signal.

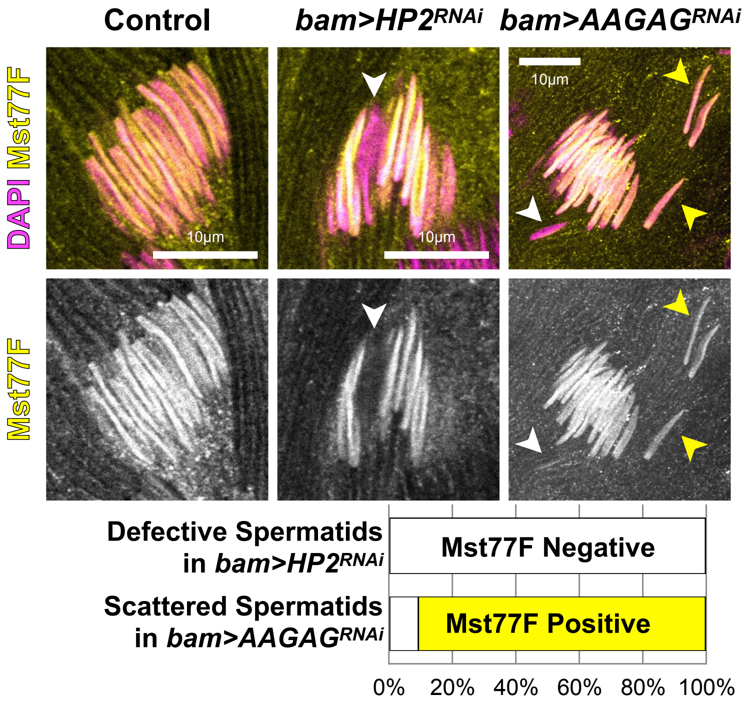


**Supplementary Fig. S5: Protamine incorporation is mostly normal in *bam>AAGAG*$^{RNAi}$**

Immunofluorescence staining of Mst77F in canoe spermatids from control, *bam>HP2*$^{RNAi}$, and *bam>AAGAG*$^{RNAi}$ animals. Frequencies of defective spermatids with (yellow arrowheads) or without (white arrowheads) Mst77F staining are shown. Only the defective spermatids (sperm DNA compaction defect in *bam>HP2*$^{RNAi}$ and sperm bundling defect in *bam>AAGAG*$^{RNAi}$) are counted (n≥36 spermatids from n≥24 testes from each genotype). Bundled spermatids in *bam>AAGAG*$^{RNAi}$ are Mst77F positive and do not exhibit a DNA compaction defect as in *bam>HP2*$^{RNAi}$. (n = 36 defective spermatids, and n = 44 scattered spermatids were counted).

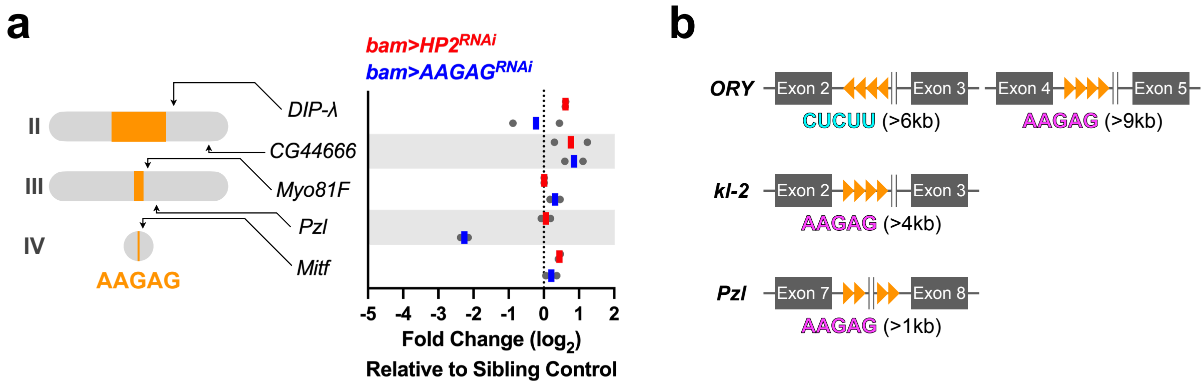


**Supplementary Fig. S6: Genomic location of AAGAG satellite DNA**

(a) Cytological map of AAGAG satellite DNA (orange) and genes with gaps in the assembly, which are potential sites for intronic AAGAG. RT-qPCR using primers to amplify the last exon of the indicated genes in *bam>HP2*$^{RNAi}$ and *bam>AAGAG*$^{RNAi,}$ compared to the control, with technical duplicates for each genotype.

(b) The schematic of gigantic introns in *ORY*, *kl*-2, and *Pzl* genes, where AAGAG satellite repeats are found next to the gaps in the genome assembly.

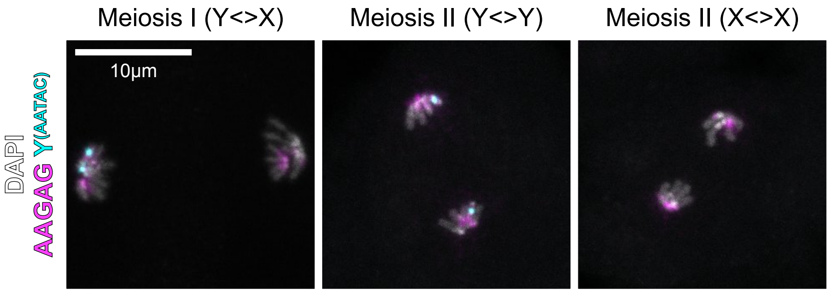


**Supplementary Fig. S7: Meiotic chromosome segregation appears normal in HP2 depletion**

DNA FISH on meiotic chromosome spread from the adult testis using AAGAG and Y-specific AATAC satellite DNA probes (n=20 testes).

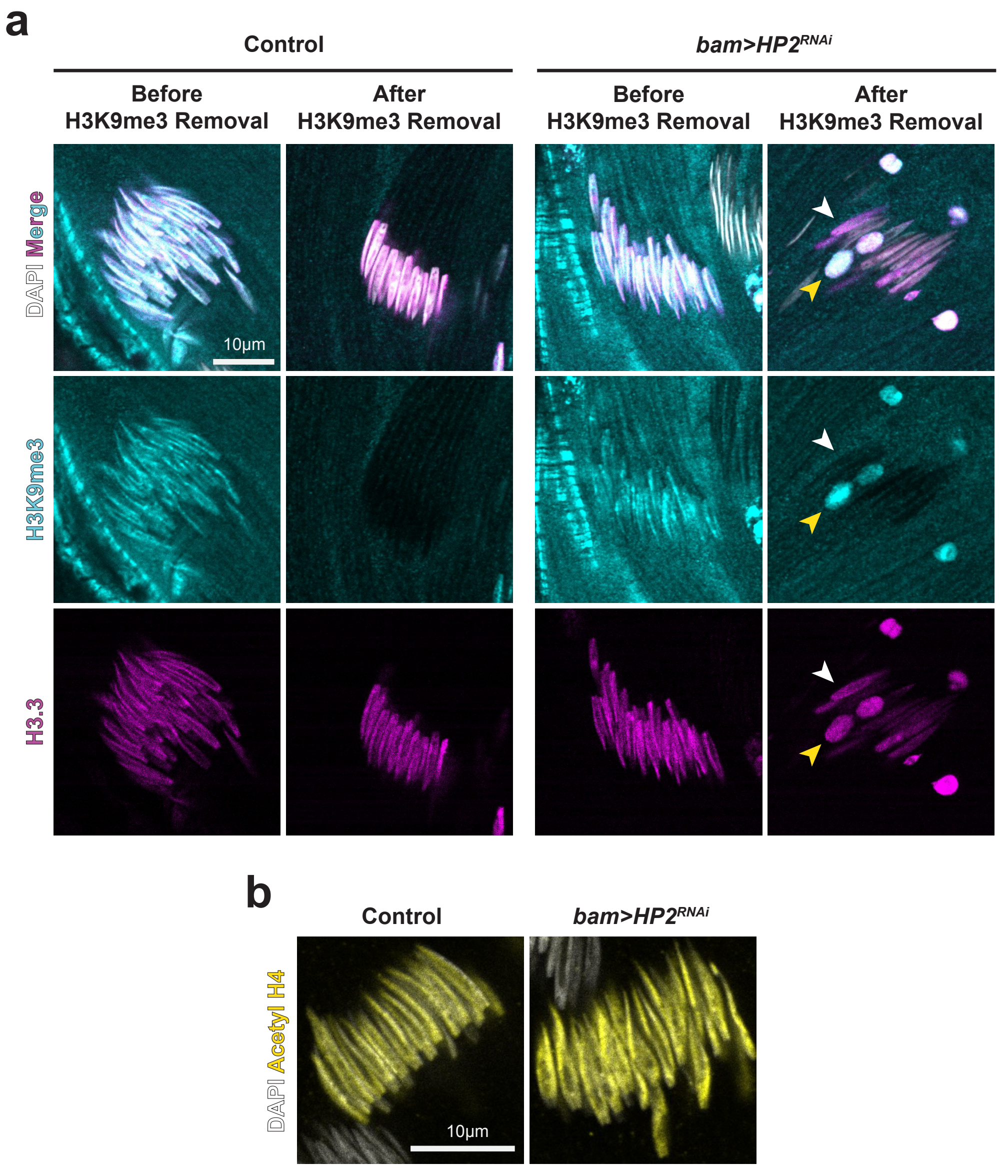


**Supplementary Fig. S8: Orderly remodeling of spermatid chromatin is compromised in *bam>HP2*$^{RNAi}$.**

(a) Immunofluorescence staining of spermatids in control vs. *bam>HP2*$^{RNAi}$ with H3K9me3 and H3.3-Dendra2 during spermatid chromatin remodeling. White arrowheads indicate spermatid nuclei that successfully removed H3K9me3, and yellow arrowheads indicate spermatid nuclei that retained H3K9me3 and failed to undergo sperm DNA compaction (n=15 testes for each genotype).

(b) Immunofluorescence staining of spermatids in control vs. *bam>HP2*$^{RNAi}$ with acetyl H4 (n= 15 testes for each genotype).

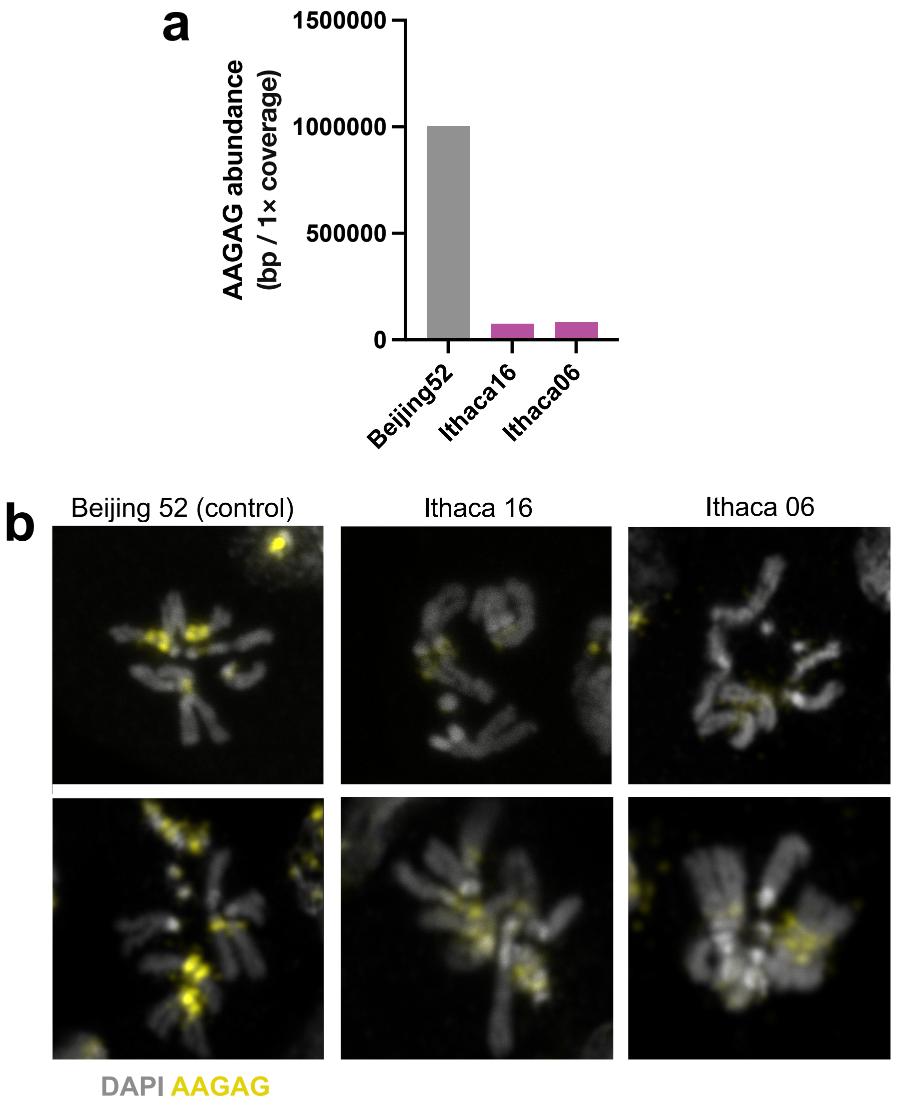


**Supplementary Fig. S9: Abundance of AAGAG satellite DNA in the D. melanogaster lines used.**

(a) AAGAG satellite DNA abundance in three *D. melanogaster* lines measured by k-Seek. Data from [1]. Beijing 52 contained the highest level of AAGAG (1,003,487 bp), whereas Ithaca 16 and Ithaca 6 lines contain much less (75,460 bp and 81,410 bp, respectively).

(b) DNA FISH on mitotic chromosome spread from the larval brain (from Beijing 52, Ithaca 16 and Ithaca 06) using the AAGAG satellite DNA probe.

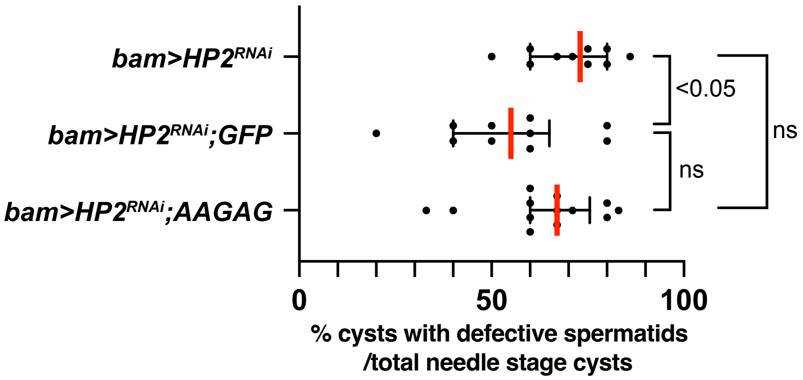


**Supplementary Fig. S10: Expression of AAGAG RNA does not rescue the *HP2*$^{RNAi}$ phenotype**

UAS-AAGAG (described in [2]) was expressed together with the *HP2*$^{RNAi}$ construct, which showed no rescue of the defective sperm DNA compaction phenotype. UAS-GFP served as a control of UAS-AAGAG transgene (to match the number of UAS transgenes driven by a single gal4 driver). Each dot represents an individual testis; red line, median; black lines, interquartile range. N = 10, 10, 13 testes were scored for *bam>HP2*$^{RNAi}$, *bam>HP2*$^{RNAi}$ *UAS-GFP*, *bam>HP2*$^{RNAi}$ *UAS-AAGAG,* respectively. The p-value was calculated by the Mann-Whitney U test (two-tailed): *bam>HP2*$^{RNAi}$ vs *bam>HP2*$^{RNAi}$ *UAS-GFP*, p = 0.0397; *bam>HP2*$^{RNAi}$ vs *bam>HP2*$^{RNAi}$ *UAS-AAGAG*, p = 0.2995; *bam>HP2*$^{RNAi}$ *UAS-GFP* vs *bam>HP2*$^{RNAi}$ *UAS-AAGAG*, p = 0.1371, and a p-value of less than 0.05 is judged as statistically significant. ns, not significant. Exact numbers of samples can be found in the Source Data file.

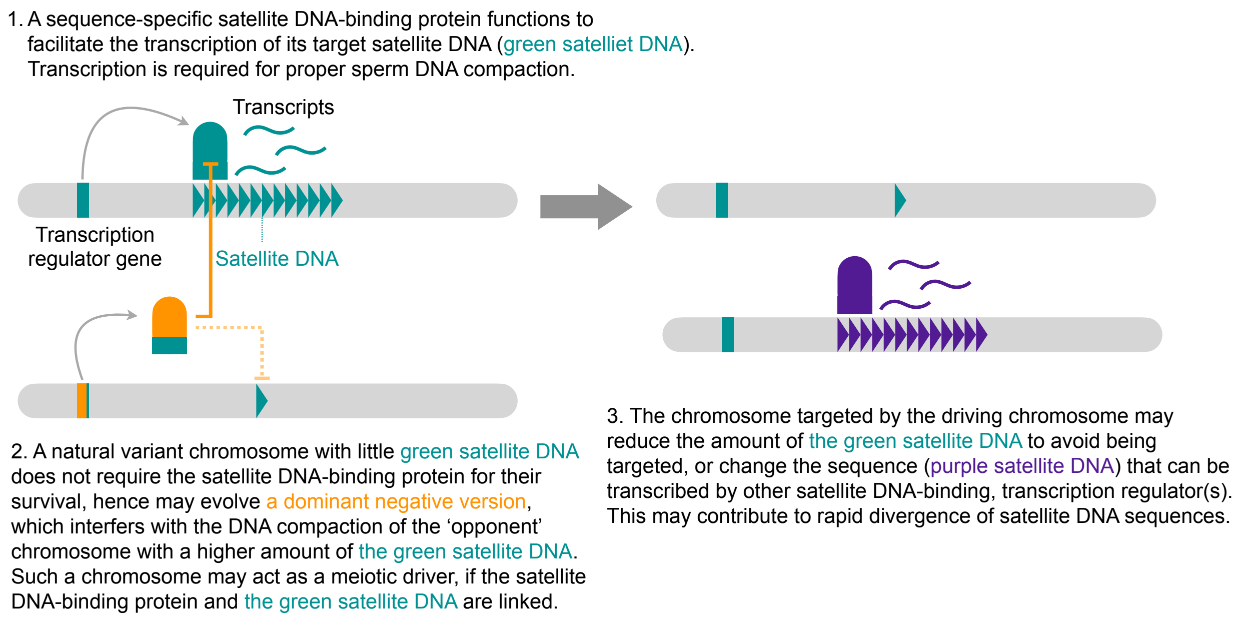


**Supplementary Fig. S11: Potential emergence of a selfish driver based on the proposed model of satellite DNA transcription and sperm DNA compaction.**

A satellite DNA-binding protein ('transcription regulator') mediates transcription of its target satellite DNA to facilitate sperm DNA packaging (1). A variant chromosome with a reduced amount of the target satellite DNA does not require the transcription regulator and may evolve a dominant-negative version of the transcription regulator, leading to preferential killing of the chromosome containing a higher amount of the target satellite DNA (2). This might lead to a meiotic drive phenomenon. The chromosome with a higher amount of the target satellite DNA may acquire resistance to the driver, either by reducing the amount of the target satellite DNA or changing the sequence, which can be transcribed by other transcription regulators. This may contribute to rapid divergence of satellite DNA (3).

**References for Supplementary Figures:**

1. Wei, K.H., Grenier, J.K., Barbash, D.A. & Clark, A.G. Correlated variation and population differentiation in satellite DNA abundance among lines of Drosophila melanogaster. *Proc Natl Acad Sci U S A* **111**, 18793-18798 (2014).
2. Mills, W.K., Lee, Y.C.G., Kochendoerfer, A.M., Dunleavy, E.M. & Karpen, G.H. RNA from a simple-tandem repeat is required for sperm maturation and male fertility in Drosophila melanogaster. *Elife* **8** (2019).
